# Patient-derived ascites reveals functional advantages of γδCAR T-cells within the ovarian cancer-like tumor microenvironment

**DOI:** 10.64898/2026.09.08.749386

**Authors:** Paula Hahn, Katerina Ntarntani, Isabella Micallef Nilsson, Emilija Gailiesaite, Emelie Foord, Michael Uhlin, Isabelle Magalhaes, Thomas Poiret

## Abstract

Chimeric antigen receptor (CAR) T-cells engineered from γδ T-cells show limited clinical benefit in solid tumors such as ovarian cancer (OVCA). Although γδ T-cells can detect malignant transformation and exert cytotoxicity, these properties confer potent preclinical efficacy but limited therapeutic activity and the basis for this discrepancy remains unclear. Here, we established an *ex-vivo* culture consisting of highly inflammatory patient-derived malignant ascites and hypoxia conditions, to simulate key obstacles encountered by CAR T-cells in the immunosuppressive OVCA tumor microenvironment (TME). Using this condition, we first demonstrated the presence of CAR-expressing γδ T-cells within conventional (conv)CAR T-cells, which displayed greater proliferative capacity than γδ_neg_ T-cells. We therefore produced γδCAR T-cells for parallel comparison with conventional (conv)CAR T-cells, evaluating differences in gene expression and functionality. Under TME-like conditions, γδCAR T-cells demonstrated lower viability but consistently outperformed convCAR T-cells, demonstrating favorable gene expression profiles, enhanced cytotoxicity, improved retention of cytokine secretion and degranulation and increased proliferative capacity. Importantly, we characterized an ascites protein profile that impacted the viability of both cell types comparably. Together, these findings reveal specific functional advantages of γδCAR T-cells over convCAR T-cells in the immunosuppressive OVCA TME-like condition while facing different cellular fitness constraints, providing insights into the use and limitations of γδCAR T-cells.

**One sentence summary:** γδ CAR T-cells retain superior functionality compared with conventional CAR T cells in an ovarian cancer-like tumor microenvironment

## INTRODUCTION

γδ T-cells, although representing a small subset of peripheral blood T-cells, possess effective antitumor functions. Notably, tumor-infiltrating γδ T-cells have been identified as the most favorable prognostic immune cell population across several cancers (*1*) and specific subsets of γδ T-cells, known to be more tissue-resident, have been directly associated with improved outcome in patients with triple-negative breast cancer and ovarian cancer (OVCA) (*2, 3*). γδ T-cells are cytolytic effector cells with distinct innate-like properties that distinguish them from conventional (conv) αβ T-cells. Expression of NK receptors enables recognition of “altered-self” signals such as MICA/B (*4, 5*), CD16 expression supports antibody-dependent cellular cytotoxicity (*6*) and their γδTCRs enable recognition of diverse stress- and transformation associated ligands without classical MHC restriction (*7, 8*). These features allow them to respond to a wide variety of antigen types in TCR-dependent and -independent manners to manage malignant transformation in both innate and adaptive approaches (*9*). γδ T-cells are divided into different subsets where Vδ2 T-cells are predominant in peripheral blood while Vδ2_neg_ T-cells (i.e. Vδ1 and Vδ3) are predominantly found in epithelial tissues. Vδ2_neg_ T-cells have been suggested to possess superior tumoricidal activity than Vδ2 T-cells (*10*) and to be crucial responders to immune checkpoint blockade (*11, 12*). The unique properties of γδ T-cells, coupled with their reduced risk to induce graft-versus-host disease (*13*), make them attractive for antitumor (engineered) cell therapy, including chimeric antigen receptor (CAR) approaches with an off-the-shelf potential (*14, 15*).

The ability of γδ T-cells to mediate broad tumor immune surveillance, remain functional in metastatic tumors and their association with favorable clinical outcomes make them promising candidates for solid tumor immunotherapy such as in OVCA (*3, 16*). However, γδ T-cells have also been shown to exert pro-tumor functions within the tumor microenvironment (TME) of OVCA. This duality may influence their therapeutic activity and underscores the need to understand how the TME shapes their function (*3, 17*).

In contrast to hematologic malignancies where tumor-associated antigens (TAAs), i.e. CD19 or BCMA, are uniformly expressed across targeted tumor cells, OVCA features large intratumoral antigen heterogeneity. In addition, OVCA presents a particularly complex landscape of spatially distinct TMEs including fluid (ascitic) and solid niches, that differ in their molecular and functional characteristics despite similar stromal cell composition (*18*). This limits the availability of ideal target antigens and necessitates the targeting of TAAs that are overexpressed in tumor cells while minimally expressed in healthy tissues. One example of a TAA in OVCA is mesothelin (MSLN), which has been targeted in multiple preclinical and clinical studies by different immunotherapy approaches, including MSLN-directed CAR T-cells (*19–21*). These challenges have motivated the exploration of γδ T-cells as an alternative CAR T cell strategy in preclinical studies of solid tumors, to leverage their intrinsic antitumor properties and potentially overcome challenges associated with antigen availability and immunosuppressive TME. γδCAR T-cells have demonstrated potent antitumor activity through complementary mechanisms, combining CAR-mediated target-cell recognition and clearance with CAR-independent recognition pathways mediated by endogenous γδTCRs and NK cell receptors, alongside robust cytokine production (*22–27*). However, despite these promising findings and the theoretical advantages of γδ T-cells over conventional (αβ) T-cells, ongoing early-phase clinical trials with γδCAR T-cells for solid tumors have shown limited therapeutic efficacy in patients with solid tumors (*15, 28, 29*). This discrepancy between preclinical promise and clinical benefit may be due to several factors: i) First, there is still limited knowledge of γδ T-cell biology and their functional inhibition/exhaustion mechanisms, especially in the TME; ii) γδ T-cells may be, similarly to αβ T-cells, limited by the physical barrier of solid tumors, therefore limiting their tumor infiltration; iii) there is not yet any consensus on the optimal γδ T-cell subset for cell therapy of solid tumors, therefore most strategies focus on subset-specific expansion (i.e Vδ1 or Vδ2) (*15*); iii) Clinical trials have been conducted by isolating and expanding γδ T-cells from peripheral blood, where they represent a rare population of 0.5-10% of T-cells, requiring extended culture to reach adequate cell numbers for clinical use, which may impact the final cell product quality (*30*); v) Last, existing *in vitro* and preclinical models to study γδ T-cells may not adequately recapitulate the TME of patients with advanced stage cancers.

Our study sought to investigate the γδCAR T-cell function in an *ex vivo* TME-like condition consisting of OVCA patient-derived ascites and hypoxia conditions. We aimed to compare gene expression profile, functionality such as cytokine production, degranulation, proliferation and cytotoxic capacity to conv T-cells using two different MSLN-directed CAR T-cells to understand the discrepancy of results between *in vitro*/preclinical and clinical trials. Furthermore, using targeted multiplex protein immunoassays, we also sought to identify an ascites protein signature that could provide insights into TME factors influencing the efficacy of (conv/γδ)CAR T-cell therapy.

## RESULTS

### γδ T-cells present within conventional CAR T-cells exhibit enhanced proliferation in the TME

We first aimed to characterize the TME of advanced OVCA to which effector T-cells would be exposed during CAR T-cell therapy. We quantified 41 protein levels in ascites collected from patients with advanced OVCA (stage III-IV) (n=36). Unbiased hierarchical clustering revealed distinct clusters, separating a distinct protein signature of ascites from healthy female control plasma (**Fig. 1A**). Ascites presented higher levels of cytokines characteristic of local immune activation and inflammation, including OSM, IL-6, TNF, IL1B, IL18, IFNγ, IL17C, IL-15 along with other factors associated with chronic stimulation and myeloid cell recruitment (IL-10, FLT3LG, CSF1/2/3) than controls (**Fig. 1B**). Ascites also contained significantly higher levels of protumoral soluble factors related to tumor growth and dissemination (TGFA and HGF), angiogenesis (VEGFA) and extracellular matrix remodeling (MMP12, IL-33) **(Fig. 1C**). Furthermore, ascites presented lower levels of a few soluble factors involved in immune and particularly T-cell function and presented higher levels of several chemokines responsible for recruitment of various immune cells (**Fig. S1A-B**).

**Figure 1.**
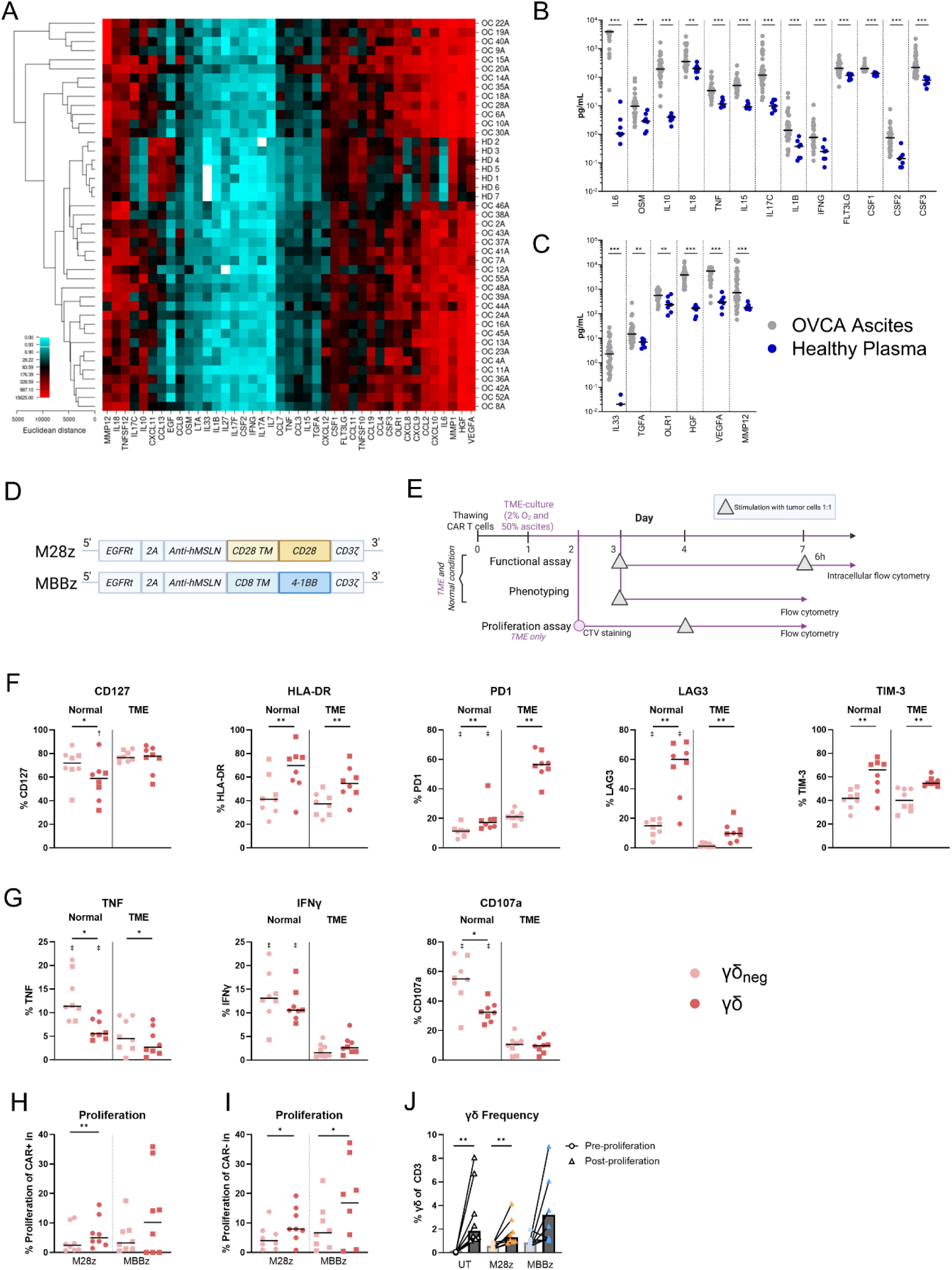
γδ T-cells present in the convCAR T cell products exhibit functional antitumor response. **(A)** Hierarchical clustering of plasma of healthy female individuals (n=7) and ascites of patients with OVCA (n=36). Heatmap was created using Euclidian distance method and the average linkage cluster algorithm. Columns show soluble factors (pg/ml); rows show individuals (HD: healthy donor, OC: ovarian cancer ascites). **(B-C)** Comparison of cytokine concentration between healthy plasma and OVCA ascites. **(D)** MSLN-CD28 (M28z) and MSLN-4-1BB (MBBz) CAR constructs. **(E**) Experimental design. Stimulations for the phenotype, functional and proliferation assays were performed with irradiated OVCAR3 M+ cells at a 1:1 E:T ratio. **F.** Frequency of CD127+, HLA-DR+, PD1+, LAG3+ and TIM3+ cells. **(G)** TNF+, IFNγ+ and CD107a+ cells in γδ T-cells and γδ-T-cells present in convCAR T-cells in normal and in TME-like conditions. **(H-I)** Proliferation of γδ+ and γδ-within convCAR T as frequency of CTV-in CAR+ (H) and CAR-(I) cells, among CD3+, γδ+ and γδ-cells. J. Frequency of γδ+ cells in UT conv T-cells and convCAR T-cells, pre- and post-proliferation assay. (F-I) n= 4, Circles represent M28z and squares represent MBBz. (H-J) n=8 in TME-like conditions. (B-C) Mann-Whitney U test with BH FDR correction across all 41 tested proteins was used to compare protein concentrations between groups. (F-J) Wilcoxon test was used for paired comparisons between cell types, constructs and conditions. Bar-free daggers represent paired comparisons between normal and TME-like conditions. */† P<0.05, **/‡ P<0.01.

To mimic this highly inflammatory TME of OVCA for functional analysis of CAR T-cells, we created an *ex vivo* condition, denoted as “TME-like”, consisting of low oxygen level (hypoxia, 2-5% O_2_) and a pool of cell-free patient-derived ascites (n=10) used at 1:1 ratio with culture medium. This *ex vivo* culture environment allowed us to evaluate CAR T cell function in TME-like conditions, using two MSLN-targeted CAR constructs, i.e. containing either a CD28 or a 4-1BB costimulatory domain (M28z and MBBz respectively, **Fig. 1D**) as previously described (*19, 20, 31–33*). As patients with decade-long remission following treatment with conventional (conv)CAR T-cells have been shown to contain a significant population of γδ T-cells (*34, 35*), we sought to investigate the function of this CAR-expressing γδ T-cell population within convCAR T cell products after 6 days exposure to the TME-like condition with target-cell stimulation (**Fig. 1E**, **Fig. S1C**). In comparison to γδ_neg_ T-cells, γδ T-cells displayed a higher proportion of HLA-DR-, PD1-, LAG3- and TIM-3-expressing cells (**Fig. 1F**). Within TME-like conditions, both T-cell subsets demonstrated a reduced frequency of LAG3+ cells compared to the normal condition, while an increase in CD127+ γδ T-cells was observed (**Fig. 1F**). γδ_neg_ T-cells demonstrated a higher frequency of TNF-expressing and degranulating cells, with no difference in IFNγ-expressing cell frequency. This function was impaired for both T-cell subsets in TME-like conditions (**Fig. 1G**). Interestingly, we observed that in TME-like conditions, the γδ T-cell population proliferated more than the γδ_neg_ T-cell population in both CAR+ (**Fig. 1H**) and CAR_neg_ (**Fig. 1I**) fractions. Accordingly, the proportion of γδ T-cells significantly increased overtime within convCAR T-cell products, which was particularly apparent within M28z-convCAR T-cells, with a similar trend for MBBz-convCAR T-cells (**Fig. 1J**). Notably, the proportion of γδ T-cells in UT conv T-cells also increased overtime (**Fig. 1J**), which reflected proliferation of γδCAR_neg_ T-cells observed in convCAR T-cell products (**Fig. 1I**).

Altogether, these results demonstrated that γδ T-cells within convCAR products showed improved proliferation than conv/γδ_neg_ CAR T-cells after tumor target-cell exposure in TME-like conditions.

### Production of γδCAR T-cells with a retained peripheral Vδ1/Vδ2 γδ T-cells proportion

To evaluate γδCAR T-cell function, we compared them to conventionally produced (conv)CAR T-cells. Briefly, γδCAR T-cells were produced from identical female donors to convCAR T-cells, by transducing pre-sorted γδ T-cells (**Fig. 2A**). This process generated two γδCAR T-cell products from as few as 0.16×10^6^ γδ T-cells yielding final γδCAR T-cell numbers ranging from 7.8 to 84×10^6^ (**Fig. 2B**) with a better expansion before (median of 9.1-fold), than after transduction (**Fig. S1D**) but was equivalent between M28z- and MBBz-γδCAR T-cells (median of 11.8-fold, **Fig. 2B**). The γδCAR T-cell products yielded a high γδ T-cell purity (>90%, **Fig. S1E and S2A**) and maintained comparable γδ T-cell subset frequencies between untransduced (UT) and the two γδCAR T-cell products (similar to peripheral blood) throughout the CAR T cell production (**Fig. 2C, Fig. S1F-H**) due to proportional increase of Vδ1 and Vδ2 γδ T-cell numbers during the production. Double negative (DN, Vδ1^−^Vδ2^−^) γδ T-cells only expanded after transduction (**Fig. S1I**). Regardless of the construct, CAR frequency was lower in γδCAR T-cells (median>40%, **Fig. S1I**) than in convCAR T-cells (median>60%, **Fig. 2D**) and no difference in CAR frequency was observed between Vδ1 and Vδ2 γδ T-cells (**Fig. 2E**). In the convCAR T-cells, a higher CD4/CD8 ratio was observed in MBBz-transduced cells compared to UT and M28z-CAR T-cells (**Fig. S1J**), as previously reported (*20, 36*).

**Figure 2.**
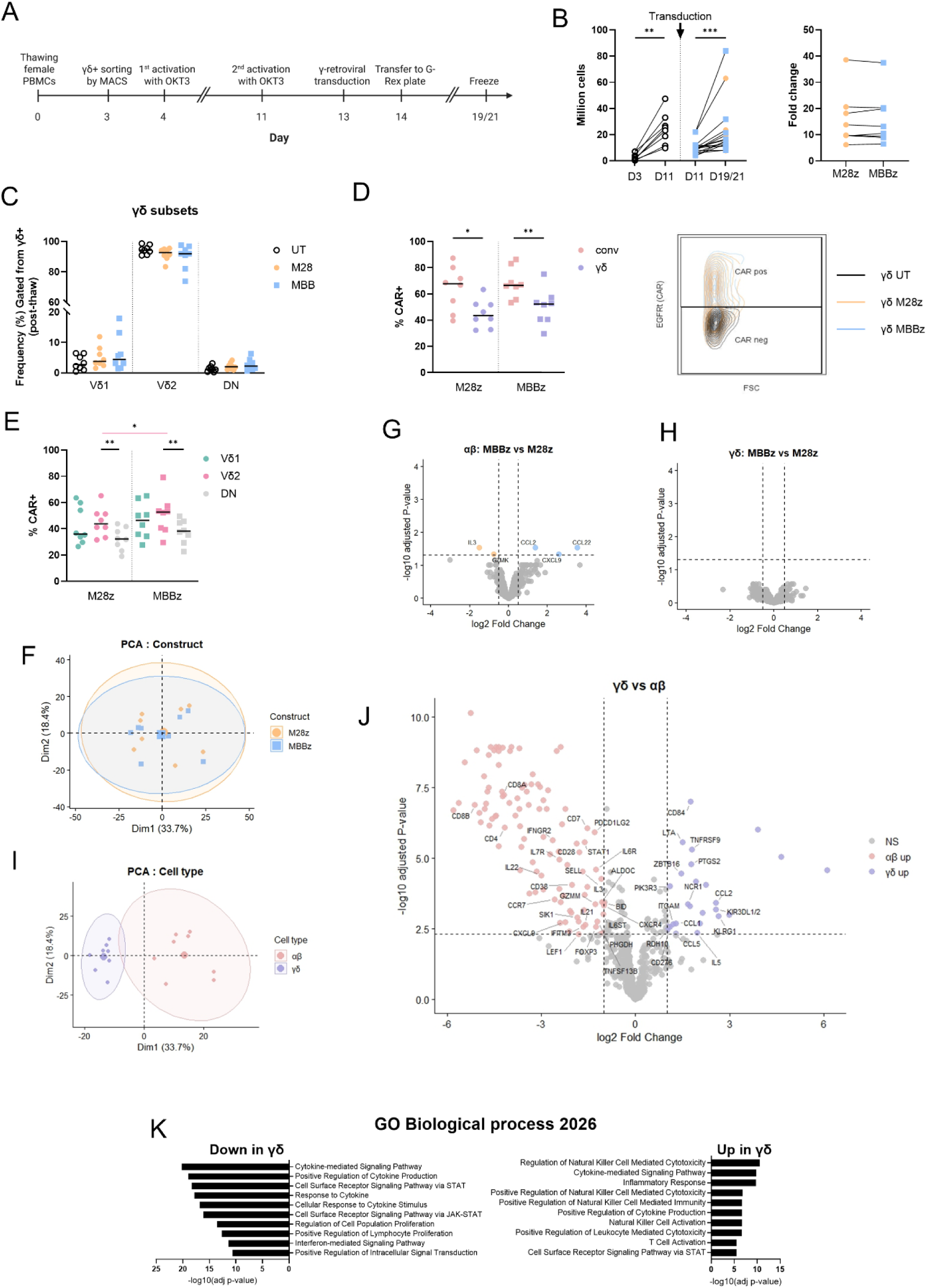
γδCAR T-cell production and transcriptional difference of γδCAR and αβCAR T-cells. **(A)** Design of the γδCAR T-cell production. **(B-C)** M28z-and MBBz-γδCAR T-cells expansion by total cell number before and after transduction and total (pre + post-transduction) fold change (FC) (B), and composition post-expansion (C)**. (D-E)** CAR (EGFRt+) frequency in convCAR (CD3+) and γδCAR (γδ+) T-cells (D) and in γδ T-cell subsets in γδCAR T-cells (E). **F.** PCA analysis of gene expression profiles comparing M28z- and MBBz-CAR T-cells. Ellipses represent the 95% confidence interval for each group. **(G-H)** Volcano plot showing differential gene expression between MBBz-and M28z-CAR T-cells within αβ (G) and γδ (H) T-cells. Differential expression was assessed using limma linear modeling with donor included as a covariate (adjusted P-value<0.05, |log2FC|>0.5). **(I)** PCA analysis of gene expression profiles comparing αβ and γδ CAR T-cells. Ellipses represent the 95% confidence interval for each group. **(J)** Volcano plot showing differential gene expression between αβCAR and γδCAR T-cells estimated using limma linear modeling with adjustment for donor and CAR construct, with labels for non-TCR genes (adjusted P-value<0.005, |log2FC|>1. **K.** Gene Ontology Biological Process enrichment analysis of genes differentially expressed between αβCAR and γδCAR T-cells (adjusted P-value<0.05, |log2FC|>0). (B-E). n=8 paired αβCAR and convCAR T-cells. (F-K). n=4 paired αβCAR and γδCAR T-cells. Wilcoxon test (B,D,E) and Friedman test with Dunn’s correction (C&E) were used for comparison. Pre-T: pre-transduction, Post-T: post-transduction. * P<0.05, ** P<0.01.

Overall, the results demonstrated a feasible production of MSLN-targeted γδCAR T-cells incorporating either a CD28 or 4-1BB costimulatory domain while preserving proportions of Vδ2 and Vδ2_neg_ T-cell subset comparable to those found in peripheral blood.

### γδCAR and αβCAR T-cells show different gene expression profiles independently of the CAR construct

To evaluate the gene expression of γδCAR T-cells, we generated four CAR T-cell products: γδ or γδ_neg_ CAR T-cells (mostly αβ T-cells, hereafter denoted as αβCAR T-cells) transduced with M28z or MBBz constructs. αβ and γδCAR T-cells were produced after separation of the γδ and γδ_neg_ T-cell subsets from expanded bulk CAR T-cells for parallel comparison using a similar production protocol. The different costimulation domains of the MSLN-targeted CAR constructs (M28z and MBBz) did not cause major differences in the CAR T-cells’ gene expression signature (**Fig. 2F**). At cell-type level, the different co-stimulatory molecules in αβCAR T-cells were associated with the relative overexpression of *IL3* and *GZMK* in M28z and the relative overexpression of *CCL2*, *CXCL9*, and *CCL22* in MBBz CAR T-cells (**Fig. 2G**), while no difference was observed for γδCAR T-cell products (**Fig. 2H**).

While the CAR T-cells did not cluster by construct or donor (**Fig. 2F, Fig. S2B**), their gene expression signatures were markedly different among T-cell subsets (γδ or αβ), with the most apparent difference caused by TCR-related genes (**Fig. 2I, Fig. S2C**). Several non-TCR genes were selectively upregulated in each CAR T-cell subset (**Fig. 2J**) highlighting distinct biological processes between the two T-cell subsets (**Fig. 2K** and **data file S4**). Relative to αβCAR T-cells, γδCAR T-cells displayed upregulation of NK cell-related pathways and downregulation of STAT-signaling and cytokine response-related pathways. Interestingly, cytokine-mediated signaling pathways were both upregulated and downregulated in γδCAR T-cells (**Fig. 2K**) in comparison to αβCAR T-cells. γδ-upregulated genes were predominantly cytokines and chemokines (e.g. *CCL1*, *CCL2*, *CCL5*, *CCL3L1*, *CCL4L1*, *TNF*, *IL5*, *IL13*, *IL12A*) while αβ-upregulated genes involved in this pathway were cytokine receptors and JAK-STAT pathway-related genes (e.g. *JAK1*, *JAK2*, *STAT1*, *STAT3*, *STAT4*, *STAT6*, *IL6R*, *IL7R*, *IFNGR1*, *IFNGR2*, **data file S4**). These results suggest that γδCAR T-cells retain a transcriptional bias toward innate effector functions and cytokine/chemokine production whereas αβCAR T-cells display greater representation of downstream cytokine signal transduction pathways.

Altogether, the costimulatory domain of the CAR constructs did not significantly impact the gene expression of αβ or γδCAR T-cell products. Importantly, γδCAR T-cells were associated with distinct biological processes from αβCAR T-cells.

### The highly inflamed, protumoral hypoxic TME impairs the γδ and conv T-cells function to different amplitudes

As γδ T-cells are able to target tumor cells by recognizing metabolic stress and/or stress-induced protein in an innate-like manner, we assessed how TME-like conditions affect the function of untransduced (UT) conv T-cells and γδ T-cells (**Fig. 3A**). In normal conditions, higher proportions of cytokine-producing and -degranulating (CD107a+) cells were found among γδ T-cells than among conv T-cells (**Fig. 3B**). TME-like conditions induced a significant decrease of function among both T-cell populations but to a stronger extent among cytokine-producing γδ T-cells (**Fig. 3C**). However, γδ T-cells maintained superior degranulation, as assessed by expression of CD107a, in TME-like conditions in comparison to conv T-cells (**Fig. 3B**). Among γδ T-cells, Vδ1 T-cells displayed a higher frequency of CD107a+ cells than Vδ2 T-cells (**Fig. 3D**). Exposure to TME-like conditions had minimal impact on the distribution of γδ T-cell subsets (**Fig. S3A**). Vδ2 T-cell functionality was markedly impaired by the TME (**Fig. 3D&E**), with a median degranulating cell frequency dropping to 1.9% whereas Vδ1 T-cells retained a median CD107a frequency of 7.4% (**Fig. 3D**). In TME-like conditions, T-cell proliferation was low (median<10%) and no differences between T-cell populations and γδ T-cell subsets were observed (**Fig. 3F&G**). After 24h co-culture with effector cells, viability of tumor cells was lower in the presence of γδ T-cells than conv T-cells at a 1:1 but not at a 1:5 E:T ratio (**Fig. 3H&I**).

**Figure 3.**
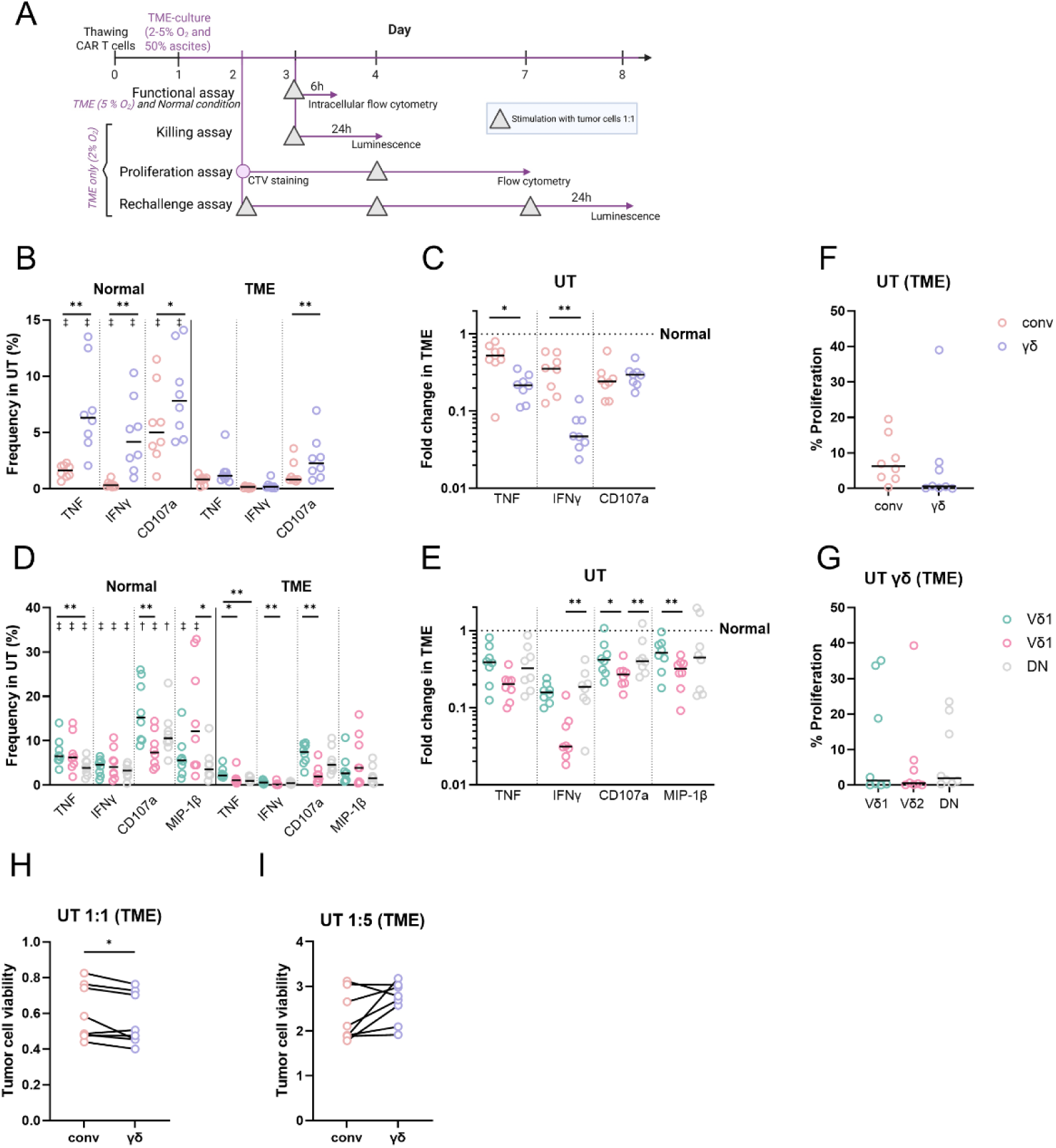
Hypoxic TME’s protein signature impairs γδ and conv T-cells. **(A)** Experimental design. Stimulation for the functional assay was performed with MSLN+ K562 cells at a 1:1 E:T ratio, and stimulation for the killing, proliferation, and rechallenge assay was performed with irradiated non-adherent MSLN+ OVCAR3+ cells at a 1:1 E:T ratio. **(B)** Frequency of TNF+, IFNγ+, CD107a+ untransduced (UT) conv and γδ T-cells in normal and TME-like conditions. **C**. Fold change induced by TME-like condition in UT conv and γδ T-cells. **(D)** Frequency of TNF+, IFNγ+, CD107a+ UT γδ T-cell subsets in normal and TME-like conditions. **(E)** Fold change induced by TME-like conditions in γδ T-cell subsets. **(F-G)** Proliferation (measured as frequency of CTV-cells) in TME-like conditions in UT conv and γδ T-cells (F) and in UT γδ T-cell subsets (G). **(H-I)** Bioluminescence of GFP/Luc+ MLSN+ OVCAR3 after 24h co-culture with effector conv and γδ T-cells in TME-like conditions at 1:1 (H), 1:5 (I) E:T ratio, normalized by control tumor cells without effector cells. (A-I) n=8, (B-F & H-I) Wilcoxon test was used for paired comparisons. B, C, F. conv were gated from CD3+ cells and γδ were gated from CD3+ γδ+ cells. Bar-free daggers represent paired comparisons between normal and TME-like conditions for each protein individually. */† P<0.05, **/‡ P<0.01.

Overall, γδ T-cells had superior functionality compared to conv T-cells in normal conditions. While TME-like conditions more strongly impacted γδ T-cell function (specifically Vδ2) than conv T-cell function, γδ T-cells retained a higher frequency of degranulating cells than conv T-cells in the TME.

### TME impacts γδCAR and αβCAR T-cells gene expression

To investigate whether the observed superior functionality of γδ T-cells was reflected on transcriptional level following CAR engineering, we compared αβ and γδCAR T-cell transcriptional responses under normal and TME-like conditions. First, as demonstrated earlier (**Fig. 2I&J**) αβCAR and γδCAR T-cell gene expression clustered separately in normal conditions (**Fig. 4A**). This clustering by cell type was less apparent under TME-like conditions but the difference between αβ and γδCAR T-cells in the TME remained driven by TCR-genes, as well as by the NK cell-like gene signature of γδ T-cells as observed in normal conditions (**Fig. S3B**). TME-like conditions impacted both constructs equally (**Fig. S3C&D**). Overall, TME-like conditions impacted αβ and γδCAR T-cells’ gene expression similarly (**Fig. 4B**, **Fig. S3E**). The genes most affected by TME-like conditions (adj P-value<0.005 and abs(FC)>1) showed a metabolic remodeling characterized by upregulation of *PDK1*, *PFKFB4*, *PGK1*, *DDIT4* and *ALDOC* consistent with adaptability of immune cells to the hypoxic TME (**Fig. 4A, Fig. S3E**). TME-exposed CAR T-cells also demonstrated upregulated genes associated with stress- and tissue-adaptation (*DDIT4*, *AREG*), important for repair and immunoregulatory T-cell programs. In addition, an elevation of *IL7R* could indicate a memory-like T-cell phenotype in the TME (**Fig. S3F**). This finding corroborates the earlier observation identifying an increased frequency of CD127+ γδ T-cells in TME-like conditions (**Fig. 1F**). Furthermore, increased expression of exhaustion-associated markers such as *TOX*, *CD38* and *CXCR4* suggested features of altered T-cell functionality which is supported by the downregulation of key cytotoxic effector (*IFNG*, *TNF*, *GZMB*, *FASLG*, *XCL1/2*) and co-stimulation genes (*TNFRSF9*, *TNFSF9*, *TNFRSF4*) (**Fig. 4A)**.

**Figure 4.**
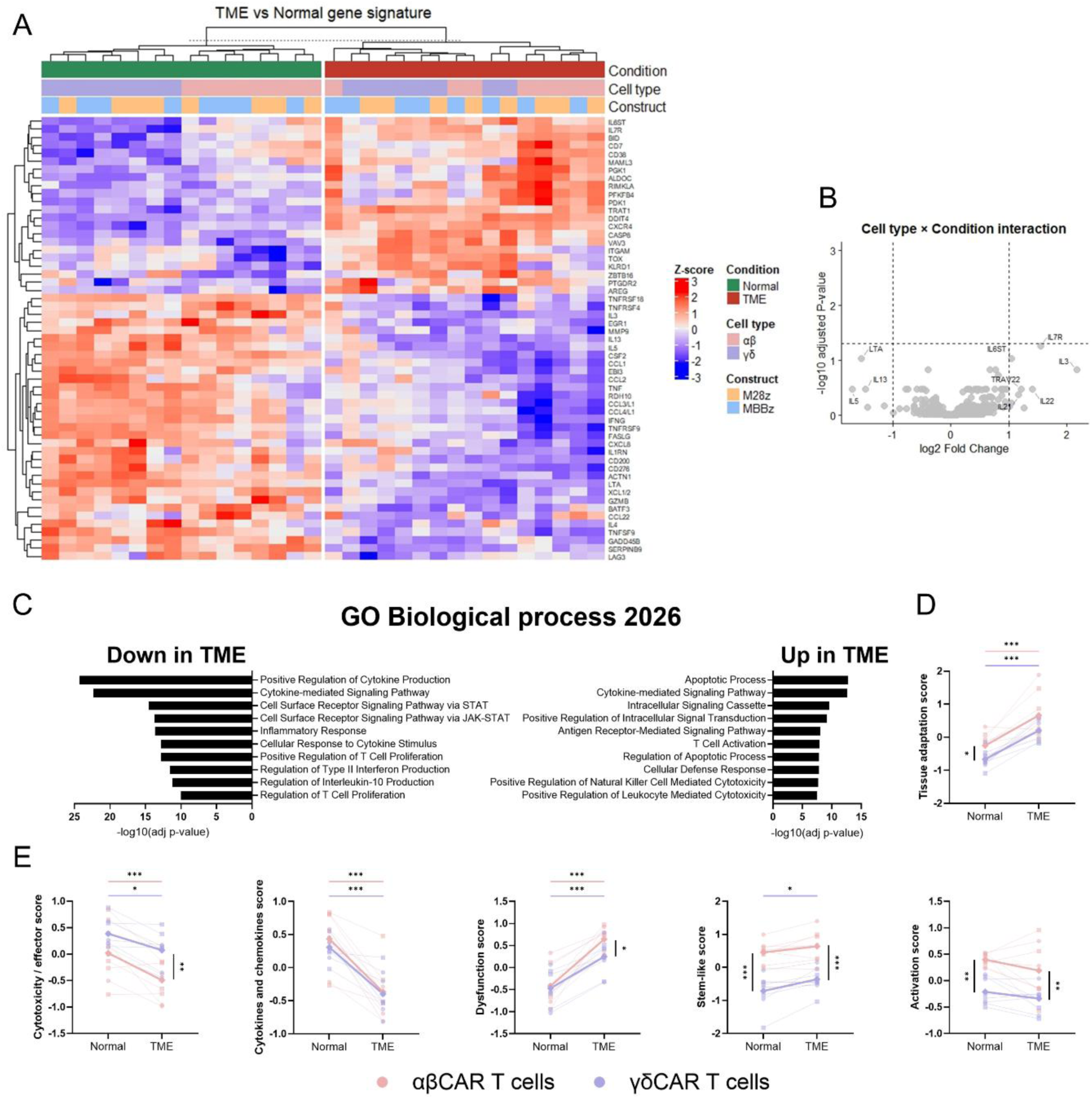
*Ex vivo* TME-like condition impacts αβCAR T-cells and γδCAR T-cells differently. **(A)** Heatmap of limma-identified differentially expressed genes between TME-like and normal conditions with adjustment for donor, cell type and construct effects. Genes were selected based on adjusted P-value<0.005 and |log2FC|>1 and visualized after gene-wise Z-score normalization followed by hierarchical clustering. **(B)** Cell type × condition interaction. Limma linear modeling was used to identify genes with differential TME responses between αβ and γδ T-cells, including donor and construct as covariates (adjusted P-value<0.05, |log2FC|>1), **(C)** GO enrichment analysis of TME-associated genes. Genes increased or decreased in TME-like relative to normal conditions (adjusted P-value<0.05) were analyzed separately using Enrichr for Biological Process 2026 enrichment. (**D-E)** Gene signature scores across normal and TME-like conditions. Signature scores were calculated as the average Z-normalized expression of the constituent genes. Thin lines connect matched donor samples between normal and TME-like conditions with αβ and γδ T-cells analyzed separately. Thick lines indicate mean signature scores, and symbols denote CAR construct identity (circles: M28z; squares: MBBz, diamonds: group means). Statistical comparisons were performed using donor-paired limma linear models adjusted for CAR construct (4 donors, 8 donor–construct paired observations per comparison). P-values were adjusted across the 14 tested modules using the BH FDR method. Significance is shown as adjusted P-values (q-values): *q<0.05, **q<0.01, ***q<0.001.

Taken together, gene expression analysis demonstrated how TME-like conditions impacted the biological pathways of (αβ/γδ)CAR T-cells with the upregulation of apoptosis, cellular defense and cytotoxicity pathways, while pathways of inflammatory response, cytokine production and STAT signaling were downregulated (**Fig. 4C and data file S4**). The TME-like conditions also induced coordinated changes across several predefined transcriptional programs (**data file S2**). Notably, both T-cell populations showed evidence of adaptation to the environment as demonstrated by an increase in the tissue adaptation score (**Fig. 4D**). However, this adaptation of T-cells also included a decrease of cytotoxicity/effector and cytokines/chemokines expression, regardless of their pro- or anti-inflammatory function and of their Th scores (**Fig. S3G&H**), resulting in an augmentation of the dysfunction score (**Fig. 4E**) without increased activation and exhaustion scores (**Fig. S3I**), probably because of the lack of antigen stimulation. Interestingly, we also observed an increase in the stemness score, especially among γδCAR T-cells, which altogether could suggest a progenitor dysfunctional state (**Fig. 4E**). While both CAR T-cell populations were impacted similarly, γδCAR T-cells retained a higher cytotoxic/effector score and lower stemness-like and dysfunction scores than αβCAR T-cells in TME-like conditions (**Fig. 4E**).

Altogether, these findings suggest a shift from a cytotoxic state (even without stimulation), towards a stress-adapted, metabolically remodeled and partially dysfunctional state of αβ and γδCAR T-cells with γδCAR T-cells retaining a higher functional score in TME-like conditions.

### γδCAR T-cells retain better function than convCAR T-cells in TME-like conditions

As TME-like conditions impacted the gene expression of γδ and αβCAR T-cells and particularly genes involved in cytotoxicity, we sought to compare γδCAR T-cell to convCAR T-cell function against MSLN-expressing tumor cells in TME-like conditions (**Fig. S4A&B**). In normal conditions, aside from a higher frequency of IFNγ+ cells in M28z-conv in comparison to MBBz-convCAR T-cells, no difference was observed between constructs across cell types (**Fig. 5A&B**). γδCAR T-cells displayed a higher proportion of cytokine-producing cells after target cells exposure compared to convCAR T-cells. Similarly to the observed transcriptional changes (**Fig. 4**), TME-like conditions significantly reduced the ability of both conv- and γδCAR T-cells to respond to antigen stimulation by cytokines or degranulation (**Fig. 5B&C**). The reduction in cytokine production was more pronounced in γδCAR T-cells than in convCAR T-cells, especially when transduced with the M28z construct, as indicated by the fold change (**Fig. 5C**). However, the proportion of TNF+ and IFNγ+ cells remained higher under TME-like conditions in MBBz-γδCAR T-cells than in convCAR T-cells (**Fig. 5B**) Interestingly, γδCAR T-cells also retained a significantly greater degranulation under TME-like conditions than convCAR T-cells (**Fig. 5B)** with degranulation being less affected by the TME **(Fig. 5C)**. These TME-related functional changes were reflected by the *TNF* and *IFNG* gene expression which decreased in TME-like conditions for both cell types and was significantly higher in γδCAR T-cells than in αβCAR T-cells (**Fig. 5D**). Gene expression of *GZMB* was reduced in TME-like conditions, in line with the observed lower proportion of CD107a+ cells, while expression of *PRF1* and *GZMH* was increased in TME compared to normal conditions for γδCAR T-cells only, which led to higher levels in γδCAR T-cells than in αβCAR T-cells in the TME (**Fig. S4C**). Consistent with an immunosuppressive TME, *FASLG* expression was also reduced in TME-like conditions in both cell types, and higher in γδCAR T-cells, following the trend of the cytotoxicity markers (**Fig. S4D**). In concordance with these observations, the frequency of triple positive (TNF+IFNγ+CD107+) cells was higher among γδCAR T-cells, especially MBBz, than in convCAR T-cells in both conditions. This triple functionality was significantly reduced by TME-like conditions (**Fig. 5E, Fig. S5A**). Looking at specific convCAR and γδCAR T-cells subsets, i.e CD4+, CD8+ T-cells or Vδ1, Vδ2 and DN γδ T-cells, TME-like conditions appeared to negatively impact their functionality similarly (**Fig. S5B-D**). In normal conditions, Vδ2 γδCAR T-cells generally showed higher functionality than other γδCAR T-cell subsets and retained this superior functionality in TME-like conditions, especially for MBBz-γδCAR T cells (**Fig. S5D**).

**Figure 5.**
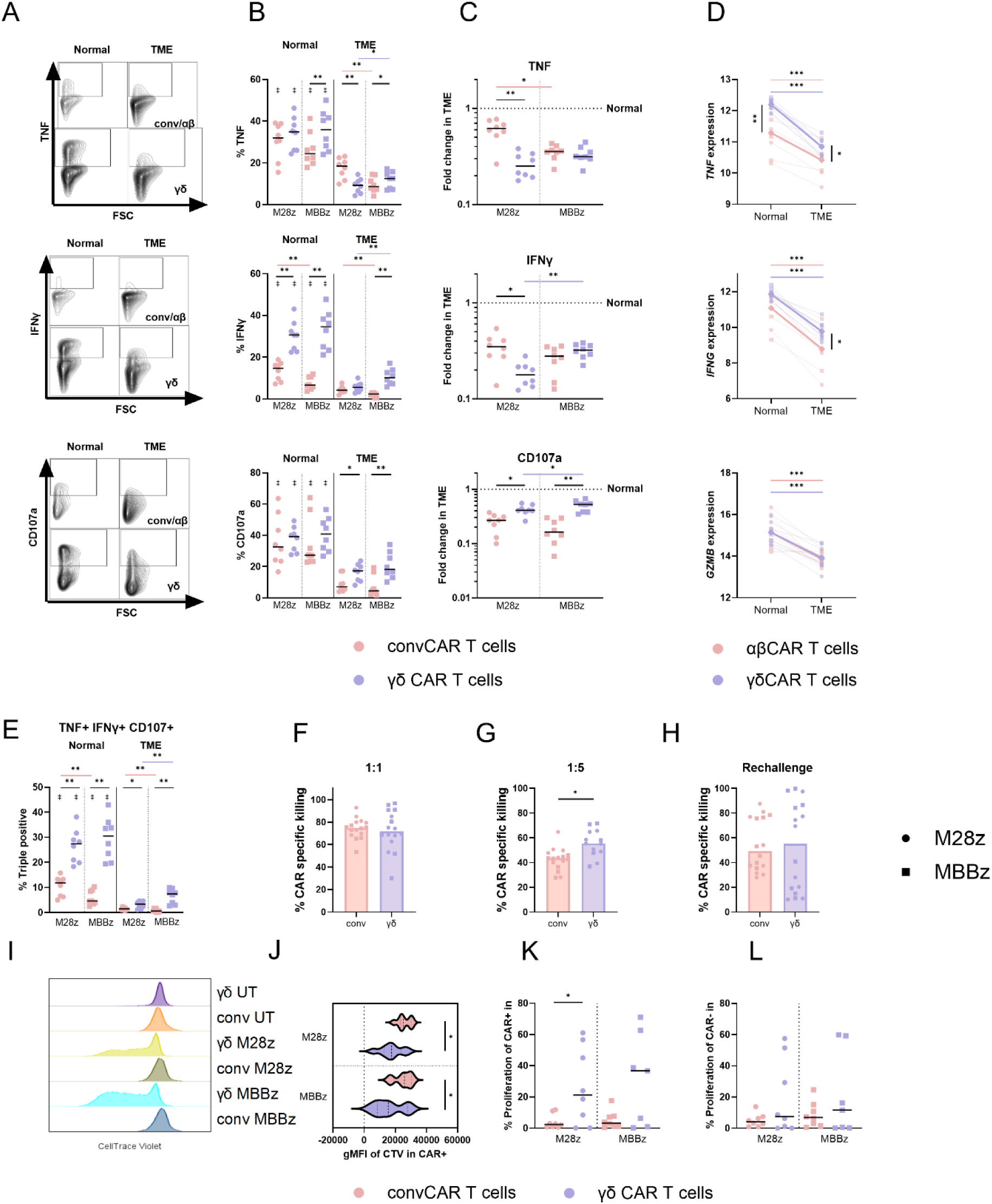
γδCAR T-cells retained better function than convCAR T-cells in TME-like conditions. **(A)** Flow cytometry plots showing TNF, IFNγ and CD107a gating in normal and in TME-like condition, for γδCAR T-cells and convCAR T-cells. **(B)** Frequency of TNF+, IFNγ+ and CD107a+ cells in γδCAR T-cells and convCAR T-cells in normal and in TME-like conditions. **(C)** Fold change of TNF+, IFNγ+ and CD107a+ frequency between TME-like and normal condition γδCAR T-cells and convCAR T-cells. **(D)** Individual gene expression across normal and TME-like conditions in αβCAR and γδCAR T-cells. Thin lines connect donor-matched samples between normal and TME-like conditions with αβ and γδ T-cells analyzed separately. Thick lines indicate mean expression values, and symbols denote construct identity (circles: M28z; squares: MBBz, diamonds: group means). **(E)** Frequency of triple-positive TNF+ IFNγ+ CD107a+ cells in normal and in TME-like conditions. **(F-G)** CAR-specific cytotoxicity of γδCAR T-cells and convCAR T-cells in TME-like conditions after 24h-stimulation with OVCAR3 cells at 1:1 (F) and 1:5 (G) E:T ratio. **(H)** CAR-specific cytotoxicity of γδCAR T-cells and convCAR T-cells in TME-like conditions after 3 repeated stimulations with OVCAR3 cells at 1:1 ratio over 6 days. Cytotoxicity is measured 24h after the third stimulation. **(I)** Flow cytometry histogram of CTV in UT and M28z- and MBBz-γδ and convCAR T-cells. **(J-L)** Proliferation of γδ and convCAR T measured as gMFI of CTV in the CTV-fraction among CAR+ cells (J) and frequency of CTV-among CAR+ (K) and CARneg (L) cells. (B, C, E-H, J-L). Wilcoxon test was used for paired comparisons between cell types, constructs and conditions. conv cells were gated from CD3+ cells and γδ cells were gated from CD3+ γδ+ cells. Bar-free daggers represent paired comparisons between normal and TME-like conditions for each protein individually. */† P<0.05, **/‡ P<0.01. (D). Statistical comparisons were performed using donor-paired limma linear models adjusted for CAR construct (4 donors, 8 donor-construct paired observations per comparison). P-values were adjusted across the 664 tested genes using the BH FDR method. Significance is shown as adjusted P-values (q-values): *q<0.05, **q<0.01, ***q<0.001.

To further assess functionality, we evaluated their killing activity in TME-like conditions. While no difference between constructs and cell types was observed at the 1:1 E:T ratio (**Fig. 5F and Fig. S5E**), γδCAR T-cells demonstrated greater cytotoxicity than convCAR T-cells at lower E:T ratio (1:5, **Fig. 5G and Fig. S5E**). After three repeated exposures with MSLN+ tumor cells in TME-like conditions, both convCAR and γδCAR T-cells were able to perform cytotoxicity towards target cells at similar frequency (median>40%, **Fig. 5H and Fig. S5E**).

Under TME-like conditions and following exposure to MSLN+ tumor cells (**Fig. 3A**), γδCAR T-cells proliferated more and at higher frequency than convCAR T-cells (**Fig. 5I-K, Fig. S6A**), but there was no difference in proliferation between γδ and conv cells within the CAR_neg_ fraction of the CAR T-cell products (**Fig. 5L, Fig. S6B&C**) suggesting that the expansion of γδCAR T-cells was driven by the CAR stimulation. No difference was observed between γδCAR T-cell subsets and CAR constructs (**Fig. S6D-F-G**).

Altogether, TME-like conditions significantly reduced the functionality of both convCAR and γδCAR T-cells, but γδCAR T-cells retained their functions more effectively, i.e. cytokine production, degranulation, cytotoxicity and proliferation compared to convCAR T-cells.

### Protein signature of ascites impact γδCAR and convCAR T-cells viability to different amplitude

We aimed to investigate if and what TME could be more deleterious to the success of CAR T-cell therapy and how specific CAR T-cell products (γδCAR and convCAR) survived when exposed to TME-associated protein profiles, i.e. ascites. First, the protein profile of ascites from individual patients with advanced OVCA was heterogenous and strongly correlated with that of paired tumor lysates (**Fig. 6A, Fig. S7A**), indicating that the liquid TME reflected the protein composition of solid TME. Interestingly, while the viability of γδCAR and convCAR T-cells after individual ascites exposure was highly correlated (**Fig. 6B**), γδCAR T-cells displayed significantly lower viability in TME-like conditions as compared to convCAR T-cells (**Fig. 6C**).

**Figure 6.**
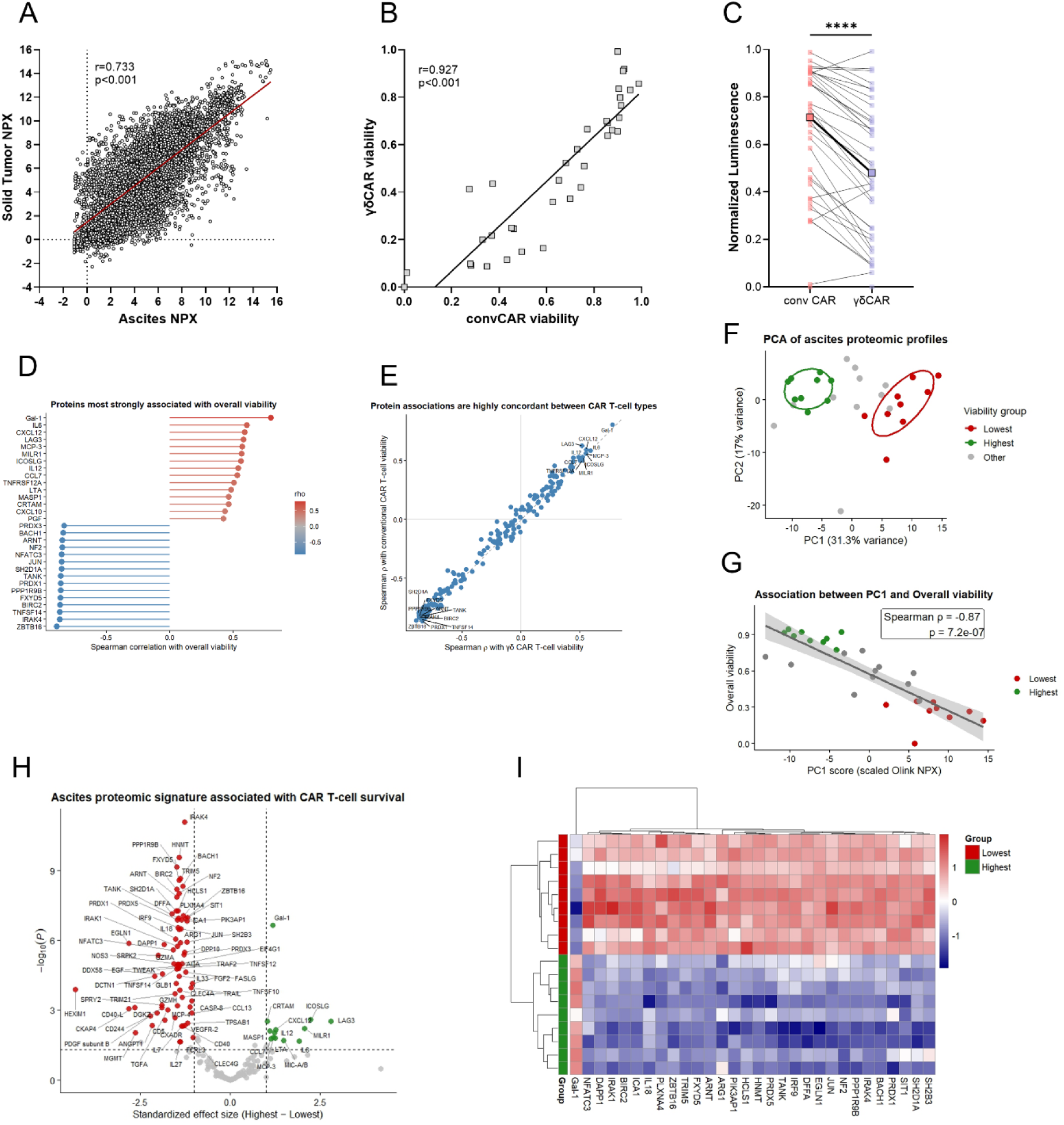
Protein signature of individual patient-derived ascites impacted the outcome of MBBz-(γδ)CAR T-cells in TME-like condition. **(A)** Correlation analysis of Normalized Protein eXpression (NPX, n=183) between paired solid tumors and ascites (n=29). (B) Correlation between γδCAR and -convCAR T-cell viability following incubation in individual patient-derived ascites placed in 2% hypoxia. Each point represents one patient sample. (C) Viability of γδCAR and -convCAR T-cells assessed by luminescence. (D) Lollipop plot showing the 15 proteins with the strongest positive and 15 strongest negative associations with overall CAR T-cell viability. Proteins are ranked by Spearman correlation coefficient (ρ), with positive correlations shown in red and negative correlations in blue. (E) Protein-level associations with γδCAR T-cell and convCAR T-cell viability. Each point represents one protein plotted according to its ρ for γδCAR T-cell viability (x-axis) and convCAR T-cell viability (y-axis). Proteins with the strongest positive and negative mean correlations are labeled. The dashed line represents the line of identity (y=x). **(F)** PCA of ascites proteomic profiles from patients with complete protein measurements (n=29). Protein abundance values were Z-score standardized across patients before PCA. The first two principal components are shown, with ellipses representing the 68% confidence region for high- and low-viability groups. **(G)** Association between PC1 score and overall CAR T-cell viability. The line represents linear regression with 95% confidence interval. **(H)** Volcano plot showing differential protein abundance between high (n=9) and low (n=9) CAR T-cell viability groups after exclusion of samples with incomplete proteomic measurements. The x-axis shows standardized effect size (difference in NPX divided by pooled median absolute deviation). Proteins with FDR<0.05 and |standardized effect size|>1 are highlighted. (**I)** Hierarchically clustered heatmap of the 30 most differentially abundant proteins between high- and low-viability groups, ranked by FDR and standardized effect size. Protein abundance values were standardized (Z-scores) across patients. Samples and proteins were clustered using correlation distance and complete linkage. Correlations were assessed using Spearman rank correlation. Differential protein abundance was assessed using linear models, with P-values adjusted for multiple testing using the BH FDR method.

Hierarchical clustering of ascites protein profiles revealed distinct patient clusters that corresponded to differences in CAR T-cell viability (**Fig. S7A**). Furthermore, the proteins most positively and negatively associated with overall CAR T-cell viability largely overlapped with those associated with γδCAR and convCAR T-cell viability analyzed separately (**Fig. 6D, Fig. S7A-C**). Indeed, protein-wise correlations with γδCAR and convCAR T-cell viability were almost identical (Spearman ρ=0.99, P<2.2×10^−16^; **Fig. 6E**), indicating that the direction and magnitude of individual protein effects were consistent across both cell types. Notably, galectin-1 (Gal-1) showed the strongest association with viability in all analyses, with a larger effect size than any other protein (**Fig. 6D&E and Fig. S7B-D**).

To identify proteins associated with CAR T-cell viability, we excluded samples with missing protein measurements and selected the nine patients with the highest and the nine with the lowest overall CAR T-cell viability. PCA of the complete protein profiles showed a clear separation between these groups (**Fig. 6F**) and PC1 was strongly negatively correlated with overall viability, convCAR and γδCAR T-cell viability (**Fig. 6G and Fig. S7E-G**). Differential protein expression analysis between the highest and lowest viability groups identified a protein signature that closely mirrored the proteins highlighted by the correlation analyses (**Fig. 6H**), further supporting their association with CAR T-cell survival. The heatmap likewise demonstrated a clear segregation of the two groups based on these differentially expressed proteins (**Fig. 6I**). The low-survival associated ascites group was characterized by increased expression of multiple proteins involved in inflammatory and immune signaling pathways including IRAK4, BIRC2, TANK, TRIM5, SHD1A and HCLS1, whereas Gal-1 was the only top-ranked protein enriched in ascites promoting the highest CAR T-cell viability.

Together, these findings demonstrate that both cell populations are impacted by similar protein signatures, that γδCAR T-cell viability is lower in TME-like conditions and that overall, a coordinated inflammatory proteomic program is associated with impaired CAR T-cell survival.

## DISCUSSION

Owing to their intrinsic antitumor activity and MHC-independent tumor recognition, γδ T-cells represent an attractive therapeutic option for solid tumors such as OVCA (*15*). However, despite promising preclinical results, γδCAR T-cell therapies have shown limited clinical benefit, and while γδ T-cells, like other infiltrating cell types, are likely affected by the immunosuppressive and hypoxic TME, the mechanisms underlying this discrepancy remain poorly understood. Here, using OVCA patient-derived ascites and hypoxia to mimic key features of the OVCA TME, we demonstrated that γδ T-cells retained equivalent antitumor responses to their γδ_neg_ counterparts across two conventional MSLN-targeting CAR T cell products. We next generated γδCAR T-cell products using the same constructs and directly compared their responses with those of conv/αβCAR T-cells under the same OVCA TME-like conditions. We found that although the TME-like conditions markedly impaired the function of both CAR T-cell products in association with important transcriptional remodeling, γδCAR T-cells retained superior degranulation, cytotoxicity, and proliferation ability. The ascites protein features associated with CAR T viability were highly concordant between both cell types, indicating that shared TME-specific constraints influence CAR T-cell fitness.

The results showed that within convCAR T-cell products, the γδ T-cell subset demonstrated an anti-tumor response with a proliferation capability outperforming αβ T-cells under TME-like conditions. This observation may suggest that, despite its low proportion in the products, at least part of the therapeutic activity of convCAR T-cells could be attributed to the γδ T-cell population present in the products. This would be in line with previous reports in long-term remission patients showing initial expansion of γδ T-cells in CAR T-cell products and their persistence up to 9 years following adoptive transfer (*34, 35*). Of interest, γδ T-cells presented a higher frequency of PD-1, especially in TME-like conditions. This observation aligns with several recent reports demonstrating that in solid tumors, γδ T-cells were highly responsive to immune checkpoint inhibitors, especially anti-PD1 blockade, and that their presence was associated with survival (*11, 12*).

This study represents a unique direct comparison of CAR T-cells containing CD28- or 4-1BB-costimulatory molecules in both γδ and conv/αβ T-cell populations. While the influence of CAR costimulatory domains has been extensively investigated in convCAR T-cells (*37, 38*), considerably less is known about the impact of these two costimulatory molecules in γδ T-cells. Similarly to others (*24*), we observed minimal differences between M28z- and MBBz-γδCAR T-cells (transcriptional profiles, proliferation, and cytotoxic activity) suggesting that γδ T-cell function may be less dependent on CAR costimulatory signals than previously observed in conv/αβ CAR T-cells (*37*). The ability of γδ T-cells to integrate CAR-independent input from their endogenous receptors such as γδTCR or other innate-like receptors remains to be investigated (*39*). This observation highlights the importance of understanding γδ T-cell biology to guide the design of next-generation CAR constructs optimized for γδ T-cells and their subsets, rather than relying on conv/αβCAR designs. Such strategies have been investigated to reduce the tonic signaling and/or improve γδ T-cell functionality (*24, 25, 40–42*).

Both Vδ1 and Vδ2 cell subsets have demonstrated cytotoxic functions against multiple tumor cell types (*43*), and Vδ1 cells are of particular interest in solid tumors because of their distinct tissue-homing, tumor-recognizing and cytotoxic properties (*44*). Yet, most previous studies of γδCAR T-cells have relied on protocols that preferentially expand Vδ2 cells using zoledronate, or Vδ1 cells using other compounds, and only few production strategies allow expansion of both subsets while preserving γδ T-cell diversity, limiting the ability to investigate γδCAR subset-specific contributions to antitumor responses (*15, 45*). For better characterization of γδ T-cell subsets and their contribution to CAR T-cell function, we generated γδCAR T-cell products that retained proportions of Vδ1, Vδ2, and Vδ1_neg_Vδ2_neg_ cells comparable to those found in peripheral blood, using zoledronate-free cultures. Using our approach, untransduced Vδ1 cells displayed greater functionality than Vδ2 cells under TME-like conditions, consistent with previous reports describing superior cytotoxicity, and stress resistance of Vδ1-enriched populations in solid tumors (*2, 10*). Unexpectedly, this was reversed following CAR transduction, despite comparable CAR expression across subsets, with Vδ2 CAR T-cells maintaining superior functionality under normal and TME-like conditions. These findings may suggest that the intrinsic antitumor properties of γδ T-cells may not necessarily translate into therapeutic benefit following CAR T-cell engineering, and that CAR signaling may differentially interact with the molecular programs of Vδ1 and Vδ2 subsets, potentially shaping their clinical performance.

Current models to investigate cell-based therapies lack patient immune factors that mimic the TME. Inspired by the method described by Tano and colleagues (*46*), we designed a clinically relevant *ex vivo* model integrating patient-derived ascites combined with hypoxia to improve clinical translation. The ascites used to reproduce specific features related to advanced OVCA including inflammatory and immunosuppressive soluble factors, enabled us to study the significant impairment of effector cells. Individual ascites protein profiles correlated with tumor protein profiles and could also be associated with cell fitness, potentially translating our finding beyond the liquid TME of OVCA. Specific protein profiles of ascites were previously associated with relapse-free survival by others (*47*). Recent work by Brown and colleagues (*48*) similarly demonstrated that OVCA ascites suppressed T and CAR T-cell functions by altering membrane lipid organization. Although their study focused primarily on metabolic mechanisms of conv T-cells and did not incorporate hypoxia, both studies converge on the functional barrier that liquid TME, i.e. ascites, represents for the success of cell-based therapies.

Despite the profound suppressive effect of TME-like conditions, γδCAR T-cells consistently displayed greater cytotoxicity, degranulation and proliferative capacity than convCAR T-cells. However, γδCAR T-cells experienced reduced viability following unstimulated exposure to TME-like conditions. Of note, a previous study reported that hypoxia in brain tumor models impairs γδ T-cell function while sparing αβ T-cells (*49*), so one could speculate that similar cell-specific impairments may also occur in our setup. Whether this reflects distinct metabolic requirements, activation-induced cell death, dependence on stimulation for survival, differential sensitivity to soluble factors present in ascites, or differential vulnerability following CAR transduction, remains to be determined. Yet, the reduced viability was balanced not only by enhanced function, but also by enhanced proliferation of γδCAR T-cells after stimulation with target cells. This observation resonated with the findings of Brown *et al.* describing that the activation state of T-cells prior to exposure to the ascites environment critically influences their subsequent function (*48*). In this context, the possible innate-like activation mechanisms of γδCAR T-cells, such as antibody-dependent-cell-mediated cytotoxicity (*25*), may allow them to maintain effector function despite suppressive environmental factors. Altogether, these findings offer clues to understanding the discrepancy between promises of *in vitro* γδ T-cell evaluation in solid tumors and limited success in the clinic (*15, 29*).

Finally, the protein analysis revealed large heterogeneity among OVCA patient-derived ascites and demonstrated that individual protein signatures largely affected CAR T-cell viability. Viability of both CAR T-cell products in individual ascites was highly correlated, indicating that the ascites’ proteomic profile generally did not favor either γδCAR or convCAR T-cells product. Nevertheless, the identification of signatures associated with high or low CAR T-cell survival may have important clinical implications, as such biomarkers could direct patient stratification and identify individuals most likely to benefit from (γδ)CAR T-cell therapy. Moreover, several of the soluble factors associated with reduced CAR T-cell persistence were involved in suppressive-inflammatory signature (ARG1, IL18, SIT1, IRAK1, IRAK4, EGLN1, SH2B3). These observations raise the possibility of a therapeutic modulation of the TME, for example through IRAK4 inhibition agents such as emavusertib (CA-4948, (*50*)), that may improve CAR T-cell fitness in patients with advanced OVCA. Unexpectedly, higher Gal-1 protein levels in ascites were associated with better CAR T-cell survival, a molecule generally indicative of bad prognosis (*51, 52*) and considered immunosuppressive within the OVCA TME and deleterious for T-cells (*53*). However, the association observed here could suggest that, within the ascites microenvironment of advanced OVCA, Gal-1 abundance may reflect a broader ascites-proteomic context that supports CAR T-cell fitness rather than directly facilitating CAR T-cell function. Rather than being driven by a single dominant mediator, CAR T-cell survival likely reflects the integrated effects of multiple interacting pathways within the TME. Future studies evaluating γδCAR T-cells in combination with TME-targeting agents may therefore represent a promising avenue for clinical translation.

Our study has several limitations; the *ex vivo* design cannot fully recapitulate the complexity of the TME, including its spatial organization, cellular heterogeneity and multicellular interactions. In addition, the availability of ascites samples limited the scope of some analyses. Despite reproducing two key TME conditions including hypoxia and cell-free ascites, our experimental setup does not allow us to distinguish their individual contributions to the effect observed on (γδ)CAR T-cells. Nevertheless, by comparing patient-derived ascites, gene expression profiling and functional analysis, we provide evidence that γδCAR T-cells retain superior antitumor activity than convCAR T-cells under conditions mimicking advanced OVCA liquid TME. These findings identify functional advantages, subset-specific contributions and TME-associated susceptibility of γδCAR T-cells and contribute to the understanding of γδ T-cells in the context of TME. Altogether, this study supports further preclinical investigations of γδCAR T-cells in a relevant OVCA model.

## MATERIALS AND METHODS

### Study design

The objective of this experimental study was to compare the functionality of γδCAR to convCAR T-cells in in-house TME-like conditions and to investigate mechanisms that may contribute to the limited clinical efficacy of γδCAR T-cell therapies. The TME-like conditions consisted of pooled advanced OVCA patient ascites mixed with culture medium and placed in hypoxia. Peripheral blood mononuclear cells (PBMCs) from healthy female donors were used in a paired manner to produce γδ and convCAR T-cells containing one of two constructs, resulting in 4 different CAR T-cell products used for downstream paired analyses by cell type, condition and construct. Targeted RNA expression profiling was performed to characterize gene expression profiles of the different CAR T-cell populations and constructs, and the effect of TME-like conditions. Functional analyses were performed using flow cytometry and luminescence based-cytotoxic assays. Multiplex protein profiling was used to characterize the protein composition of OVCA patients’ ascites and tumors. Sample sizes were defined at study initiation based on the expected availability of ascites material. Assessment of CAR T-cell viability using individual ascites samples was incorporated during the study after identification of interpatient variability in ascites protein profiles. Detailed descriptions of experimental designs, sample allocation, and data analyses are provided in the materials & methods section and figure legends. Information regarding antibodies, reagents and software used in the study are reported in the below sections and in **Supplementary Materials and Table S1**. All individual-level data are available in **data file S1.**

### Healthy donor and Patient Sample collection

PBMCs were isolated from buffy coats obtained from healthy female donors at Karolinska University Hospital, Sweden (Dnr 2025-04328-01) using Ficoll-Paque density gradient separation (GE Healthcare). PBMCs were frozen in fetal bovine serum (FBS) and 10% DMSO. Ascites (n=36) and tumors (n=29) were collected during primary tumor debulking for advanced epithelial OVCA (FIGO>II) without pretreatment at the Karolinska University Hospital (Solna, Sweden) after obtaining written patient consent as previously described (*3*) and frozen at –80°C. The study was in accordance with the Declaration of Helsinki and approved by the Regional Ethical Review Board of Stockholm, Sweden (2013/2161-31/2, 2016/1136-32, 2016/1631-32, and 2020-03761).

### TME-like conditions preparation

For functional assays, different conditions were established: “Normal condition” was regular media free of ascites and an oxygen level set at 21%. “TME-like conditions” were a culture media consisting of pooled cell-free ascites (n=10) used at 1:1 ratio with regular culture media and low oxygen level (2-5% O_2_). Regular culture medium for convCAR T-cells was complete AIM-V medium (Gibco) supplemented with 5% heat-inactivated Human AB Serum (Sigma-Aldrich, USA) and 300 IU/mL IL-2 (Proleukin; Novartis). Regular culture medium for γδCAR T-cells was complete CTS™ OpTmizer™ T-cell Expansion medium (Gibco, USA) medium supplemented with 5% heat-inactivated Human AB Serum and 70 ng/mL IL-15 (premium grade, Miltenyi Biotec, Germany).

### Data analysis and statistical analysis

GraphPad Prism 10 (GraphPad Software, San Diego, CA, USA), bioinformatics tool Online CIMminer, and Rstudio were used for figure generation and statistical analyses. Analysis was performed using two-tailed nonparametric tests: Wilcoxon test and Friedman test with Dunn’s correction for paired samples (conditions, subsets, constructs), and Mann-Whitney test for unpaired samples (Healthy individuals vs. OVCA patients). For gene and multiplex protein analyses, P-values were adjusted using the Benjamini-Hochberg (BH) method to control the false discovery rate (FDR). Correlation was assessed using Spearman’s rank correlation. Threshold significance was set at P<0.05. Corresponding P-values and applied statistical tests are specified in the figure legends. Unless stated otherwise, median values are shown as horizontal lines. All individual-level data for n<20 are presented **in data file S1**. Details on analysis of protein and gene expression can be found in **data files S2-4**.

## Acknowledgements

We thank all patients enrolled in this study. We would also like to thank Charlotte Klynning and all clinicians and nurses involved in the collection of tumor and ascites, Prof. M. Sadelain, Memorial Sloan Kettering Cancer Center (MSKCC), New York, USA, for donating the different CAR constructs used in this study, Arwen Stikvoort for her assistance with sample preparation for Olink analyses, and Emilia Bogdanovski for her assistance with the CAR T-cell viability assessment. We also acknowledge the contributions of the KIGene Core Facility (Karolinska Institutet, Sweden) for assistance with Nanostring experiments and the MedH Flow Cytometry Core Facility financed by the Infrastructure Board at Karolinska Institutet for providing instruments for cell sorting and technical expertise.

## Funding

The work was supported by grants from

Center for Innovative Medicine (FoUI-1023424) to TP,

Karolinska Institutet Research Grant (2024–02323) to TP,

Swedish Research Council (2021–01755) to MU,

Swedish Cancer Society (22 2257Pj) to MU,

Center for Innovative Medicine (FoUI-985704) to IM,

Swedish Cancer Society (24 3496 Pj) to IM,

Cancer Research Foundations of Radiumhemmet (251353) to IM.

## Author contributions

TP developed the concept. PH, IM and TP designed the study. PH, AN, IMN, EG and TP performed the experiments. MU planned and conceptualized the samples collection, EF collected and processed patient samples. MU and PH planned and conceptualized the protein analysis, PH and TP performed all data analysis and visualization. PH and TP supervised the study. TP, IM and MU acquired funding. PH and TP wrote the manuscript. All authors reviewed and approved the final manuscript.

## Data and materials availability

All data associated with this study are in the paper or the Supplementary Materials.

## Supplementary Materials

### Supplementary Materials and Methods

#### Cell lines

K562 cells were cultured in RPMI-1640 (Hyclone) supplemented with 10% FBS (Hyclone) and 1% penicillin-streptomycin (Gibco). OVCAR-3 was cultured in RPMI-1640 supplemented with 20% FBS, 1% penicillin-streptomycin and 0.01 mg/mL insulin (Sigma-Aldrich). Tumor cells were transduced with γ-retroviral vectors encoding for human MSLN with or without green fluorescent protein (GFP)/firefly luciferase fusion protein and sorted as previously described (*20*), and named K562 M+, K562 M+G+, OVCAR3 M+ and OVCAR-3 M+G+. The sorting was performed at the MedH Flow Cytometry Core Facility that receives funding from the Infrastructure Board at Karolinska Institutet.

#### CAR T-cell production

For γδCAR T-cell production, PBMCs were thawed and rested in complete γδ T-cell medium consisting of CTS™ OpTmizer™ T-cell Expansion medium (Gibco, USA) supplemented with 5% heat-inactivated Human AB Serum, (Sigma-Aldrich, USA) 1x Glutamax™ Supplement (Gibco, 1% Penicillin/Streptomycin (Gibco) and 70 ng/mL human IL-15 (premium grade, Miltenyi Biotec, Germany). γδ T-cells were isolated using the Anti-TCRγ/δ MicroBead Kit (Miltenyi Biotec) and cultured in a G-REX plate (24-wells, Wilson Wolf, USA) overnight before activation with 50 ng/mL CD3 monoclonal antibody (OKT3; Biolegend, USA). After 1 week, expanded γδ T-cells were activated again with anti-CD3 antibody 48h before MSLN-CAR transduction as previously described (*33*). The CAR constructs contain either a CD28-CD3ζ (M28z) or a 4–1BB-CD3ζ (MBBz) co-stimulatory domain (*20*). After transduction, γδCAR T-cells were transferred to a G-REX plate and expanded for 5-7 days. All expansions were conducted at 37°C and 5% CO2, and complete CTS medium or only cytokine content was replenished every 2-3 days. Cell content was assessed weekly by flow cytometry.

For conventional (conv)CAR T-cell production, PBMCs were thawed and rested in complete AIM-V medium (Gibco) supplemented with 5% Human AB Serum (Sigma-Aldrich, USA) and 300 IU/mL IL-2 (Proleukin; Novartis). Activation and transduction were performed in complete AIM-V medium as described previously (*19, 33*).

#### Characterization of γδ T-cells in conventional CAR T-cell products

ConvCAR T-cells and UT cells were thawed and rested overnight before separated culture at 1×10^6^/mL in TME-like or normal condition for 48 hours. CAR T-cells were then stimulated at a 1:1 ratio with irradiated (55Gy) OVCAR-3 M+ cells adjusted to the truncated EGFR (EGFRt) frequency for 96 hours without cytokines. EGFRt served as a non-immunogenic reporter gene for CAR expression. Cells were then analyzed for phenotype (**Suppl Table 1**) and functionality such as degranulation and cytokine production as described below.

#### Proliferation assay

Cells were thawed and rested overnight before 24h conditioning in TME-like conditions. CAR T-cells and UT cells were then stained with Cell Trace Violet (Invitrogen) at a concentration of 0.05% in PBS according to manufacturer instructions. Following 48 hours in TME-like conditions, cells were stimulated at a 1:1 E:T ratio with irradiated (55Gy) OVCAR-3 M+ cells for 72 hours in TME-like conditions without cytokines. For the assay, the CAR T-cell number was adjusted within each donor based on the transduction rate (EGFRt frequency) to have equal transduction rates between constructs and cell types. Cells were then stained with surface antibodies before flow cytometric analysis.

#### Cell preparation for gene expression analysis

For gene expression analysis, PBMCs (n=4) were thawed and rested overnight in complete γδ T-cell medium, activated and transduced as described above for γδCAR T-cell production. γδ T-cells were then sorted from the bulk CAR T-cell products using the Anti-TCRγ/δ MicroBead Kit, and expanded separately from αβCAR T-cells, in complete γδ T-cell medium in G-Rex plates at 37°C and 5% CO2 with medium and/or IL-15 replenishment every 2-3 days for two weeks. γδCAR T-cells and αβCAR T-cells were then cultured separately at 1×10^6^/mL in complete γδ T-cell medium + 50% patient-derived ascites pool (n=10) in 5% hypoxia or in 100% complete γδ T-cell medium in normoxia for 48 hours.

CAR T-cells were then enriched using MACS MS columns following staining with a human EGFR biotinylated antibody (R&D systems) and anti-biotin microbeads (130-090-485, Miltenyi biotec). 100 000 CAR T-cells were pelleted and lysed by vortexing for 1 min in RLT buffer (at 10 000 cells/ul) (Qiagen, Germany), and frozen at −80°C before analysis. 10,000 cells were analyzed for gene expression using the nCounter CAR-T Characterization panel (Nanostring Technologies) at the Nanostring MAX/FLEX platform (KIGene core facility, Karolinska Institutet, Stockholm, Sweden, **data file S2**).

#### Flow cytometry

For extracellular staining, cells were washed, stained with antibodies in **Suppl Table 1** for 30 min at 4°C in FACS buffer (PBS with 1% FBS), washed, and resuspended in FACS buffer for acquisition. All experiments were run on a CytoFLEX (Beckman Coulter, USA) and analyzed in FlowJo v.10.10.0 (BD).

#### Degranulation and cytokine production analysis

γδCAR T-cells, convCAR T-cells and matched control (donor untransduced, UT) cells were assessed by FACS and cultured separately at 1×10^6^/mL in TME-like or normal conditions for 48 hours. Cells were then stimulated with K562 M+ cells at a 1:1 effector:target (E:T) ratio. For the assay comparing γδCAR T-cells to convCAR T-cells (**Fig. 5**), the CAR T-cell number was adjusted within each donor based on the transduction rate (EGFRt frequency) to have equal transduction rates between constructs and cell types. The stimulation was performed for 6 hours in TME-like or normal conditions in the absence of cytokines. Cytokine production and degranulation were assessed by intracellular staining as previously described (*32*) using the surface and intracellular antibodies described in **Suppl Table 1**. Cells stimulated with PMA and ionomycin were used as positive control.

#### Cytotoxicity assay

CAR T-cells and UT cells were thawed and conditioned in respective TME-like conditions at 1×10^6^/mL for 48 hours as described above and stimulated at a 1:1 and 1:5 E:T ratio with irradiated (55Gy) OVCAR-3 M+G+ cells for 24 hours in triplicates in the same conditions, without cytokines. For the assay, the CAR T-cell number was adjusted within each donor based on the transduction rate (EGFRt frequency) to have equal transduction rates between constructs and cell types. Tumor cell viability was assessed by firefly luciferase gene expression using the ONE-Glo EX Luciferase Assay System reagent (Promega) according to manufacturer’s instructions, and luminescence was recorded using a CLARIOstar Multireader (BMG Labtech). Cytotoxicity was calculated as follows:

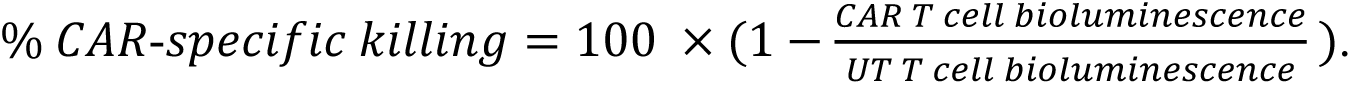

#### Rechallenge assay

CAR T-cells and UT cells were thawed and conditioned for 24 hours in respective TME-like conditions at 1×10^6^/mL. Cells were then stimulated at a 1:1 E:T ratio with irradiated (55Gy) OVCAR-3 M+ cells at timepoint 0 and after 48 hours, and with OVCAR-3 M+G+ cells after 120 hours in TME-like conditions, without cytokines. For the assay, the CAR T-cell number was adjusted within each donor based on the transduction rate (EGFRt frequency) to have equal transduction rates between constructs and cell types. After the third stimulation, cells were split into triplicates and incubated for 24h before evaluating the tumor cell viability by firefly luciferase gene expression. Cytotoxicity was calculated as the cytotoxicity assay.

#### Ascites Protein signature and Viability assessment of CAR T-cells in individual ascites

Protein signatures of patients’ ascites were analyzed using Olink Target 48 Cytokine (n=36), Target 96 Immuno-Oncology and Target 96 Immune Response (n=29;Olink Proteomics AB, Uppsala, Sweden, **data file S3**). MBBz-γδCAR and MBBz-convCAR T-cells were produced using PBMCs from 3 heathy female donors as described above. In 96-wells cell culture plate (Corning, US), 30k cells/well were seeded and cultured in 100uL cytokine-free media containing 50% patient-derived ascites for 48h in 2% oxygen. The assay was performed in duplicate screening of 36 different patients-derived ascites. After TME-like culture, cells were spun down and resuspended in 100ul RPMI supplemented with 10% FBS media. γδCAR and convCAR T-cell viability was assessed using CellTiter-Glo® Luminescent cell Viability Assay (Promega, US) according to the manufacturer’s instructions and luminescence was recorded using a CLARIOstar Multireader. Data were then transformed using Min-Max normalization before analysis.

##### Analysis of protein expression profiles

The two-dimensional clustered image map of soluble protein concentrations found in ascites of OVCA patients (n=36) and plasma of healthy female individuals (n=7) was created using the Euclidian distance method and the average linkage cluster algorithm on the bioinformatics tool Online CIMminer. Quantitative protein measurements in pg/mL from the Olink target 48 panel were included. (**Fig. 2**). Comparison of protein levels in plasma from healthy individuals and ascites of OVCA patients was performed using Mann-Whitney before using false discovery rate (FDR) by Benjamini-Hochberg (BH) method.

Prior to analysis of the associations between ascites protein abundance and CAR T-cell viability (**Fig. 6**), duplicated protein entries were removed from the dataset. Associations were assessed using Spearman rank correlation analysis based on available paired measurements (n=29–36). Correlations were calculated separately for γδCAR and conventional CAR T-cell viability, with P-values adjusted using the BH FDR method. Concordance between CAR T-cell products was assessed by correlating protein-level Spearman coefficients.

Overall CAR T-cell viability was calculated as the mean of γδCAR and convCAR T-cell viability. Patients were ranked by overall viability, and differential protein abundance between the nine highest- and lowest-viability patients was assessed using linear models after excluding samples with missing proteomic measurements (n=29). FDR correction was applied for multiple testing. PCA was performed on scaled protein abundance values from complete proteomic profiles (n=29), and associations between principal components and viability parameters were evaluated using Spearman correlation.

##### Analysis of gene expression profiles

Gene expression profiling was performed using the NanoString platform. Data were processed and normalized using the Rosalind Bioinformatics platform according to the manufacturer’s recommended workflow, (https://www.rosalind.bio/en/knowledge/normalization-methods-for-nanostring-gene-expression-from-rcc-files) before downstream analyses in Rstudio.

Gene expression module scores were generated from predefined gene sets (**data file S2**). Expression values were Z-score normalized across samples for each gene, and module scores were calculated as the mean normalized expression of all available genes within each score. No gene weighting was applied. Principal component analysis (PCA) was performed on Z-score normalized expression data within the normal (N) condition using all genes. Differential gene expression analyses were performed using the limma package to compare conditions, cell types, CAR constructs, and interaction effects. Linear models included donor and CAR construct as a covariate where appropriate. For heatmap visualization, genes were Z-score normalized across samples and displayed using hierarchical clustering. Gene ontology enrichment analysis was performed using Enrichr with the Biological Process 2026 database using selected differentially expressed gene lists (**data file S4**). All comparisons were performed using donor-matched samples with multiple testing correction performed using the BH FDR method.

### Supplementary Tables and Figure legends

**Table S1.** Antibodies and dyes used for flow cytometry.

| Label | Marker / Dye | Catalogue number | Clone | Vendor | Experiment used |
| --- | --- | --- | --- | --- | --- |
| FITC | CD4 | 555346 | RPA-T4 | BD Biosciences | ICS |
| FITC | PD1 (CD279) | 557860 | MIH4 | BD Biosciences | Phenotyping |
| FITC | TCR_Vδ1 | TCR2730 | TS8.2 | Invitrogen | ICS, Proliferation, Production D3, D11 D17, Post-thaw |
| Biotin | EGFR | FAB9577 B | Hu1 | R&D Systems | Proliferation, Production D17, Post-thaw |
| PE | Streptavidin | 405204 | - | Biolegend | Proliferation, Production D17, Post-thaw |
| PE | CD107a | 555801 | H4A3 | BD Biosciences | ICS |
| PE | TCR_$\gamma\delta$ | 130-113-512 | REA591 | Miltenyi Biotec | Phenotyping |
| PE-eFluor610 | TNF-α | 61-734942 | MAb11 | Invitrogen | ICS |
| PerCP | CD3 | 300428 | UCHT1 | Biolegend | Production D3, D11, D17, Post-thaw |
| PECy7 | CD3 | 300316 | HIT3a | Biolegend | ICS, Proliferation |
| APC | IFN-γ | 551385 | 4S.B3 | BD Biosciences | ICS |
| APC-Alexa Fluor 700 | CD127 | A71116 | R34.34 | Beckman Coulter | Phenotyping |
| APC | TCR_Vδ2 | 130-121-339 | 123R3 | Miltenyi Biotec | Proliferation, Production D3, D11, D17, Post-thaw |
| Alexa Fluor 700 | CD4 | 557922 | RPA-T4 | BD Biosciences | Production D3, D11, D17, Post-thaw |
| Alexa Fluor 700 | MIP-1b | 561278 | D21-1351 | BD Biosciences | ICS |
| Alexa Fluor 700 | IL-2 | 500320 | MAQ1-17H12 | Biolegend | ICS |
| APC-Cy7 | CD8 | 557834 | SK1 | BD Biosciences | ICS, Production D3, D11, D17, Post-thaw |
| APC-H7 | HLA-DR | 561358 | G46-6 | BD Biosciences | Phenotyping |
| VioBlue | TCR_Vδ2 | 130-101-157 | 123R3 | Miltenyi Biotec | ICS, Phenotyping |
| BV510 | CD3 | 300448 | UCHT1 | Biolegend | Phenotyping |
| BV650 | LAG-3 | 369316 | 11C3C6 5 | Biolegend | Phenotyping |
| BV650 | TCR_$\gamma\delta$ | 569510 | 11F2 | BD Biosciences | ICS, Proliferation, Production D3, D11 D17, Post-thaw |
| BV785 | TIM-3 (CD366) | 345032 | F38-2E2 | Biolegend | Phenotyping |
| Aqua | LIVE/DEAD | L34957 | - | Invitrogen | ICS, Production D3, D11, D17, Post-thaw |
|  | CellTrace Violet | C34557 | - | Invitrogen | Proliferation |
|  | 7AAD | 51-68981E | - | BD Biosciences | Proliferation |

**Figure S1.**
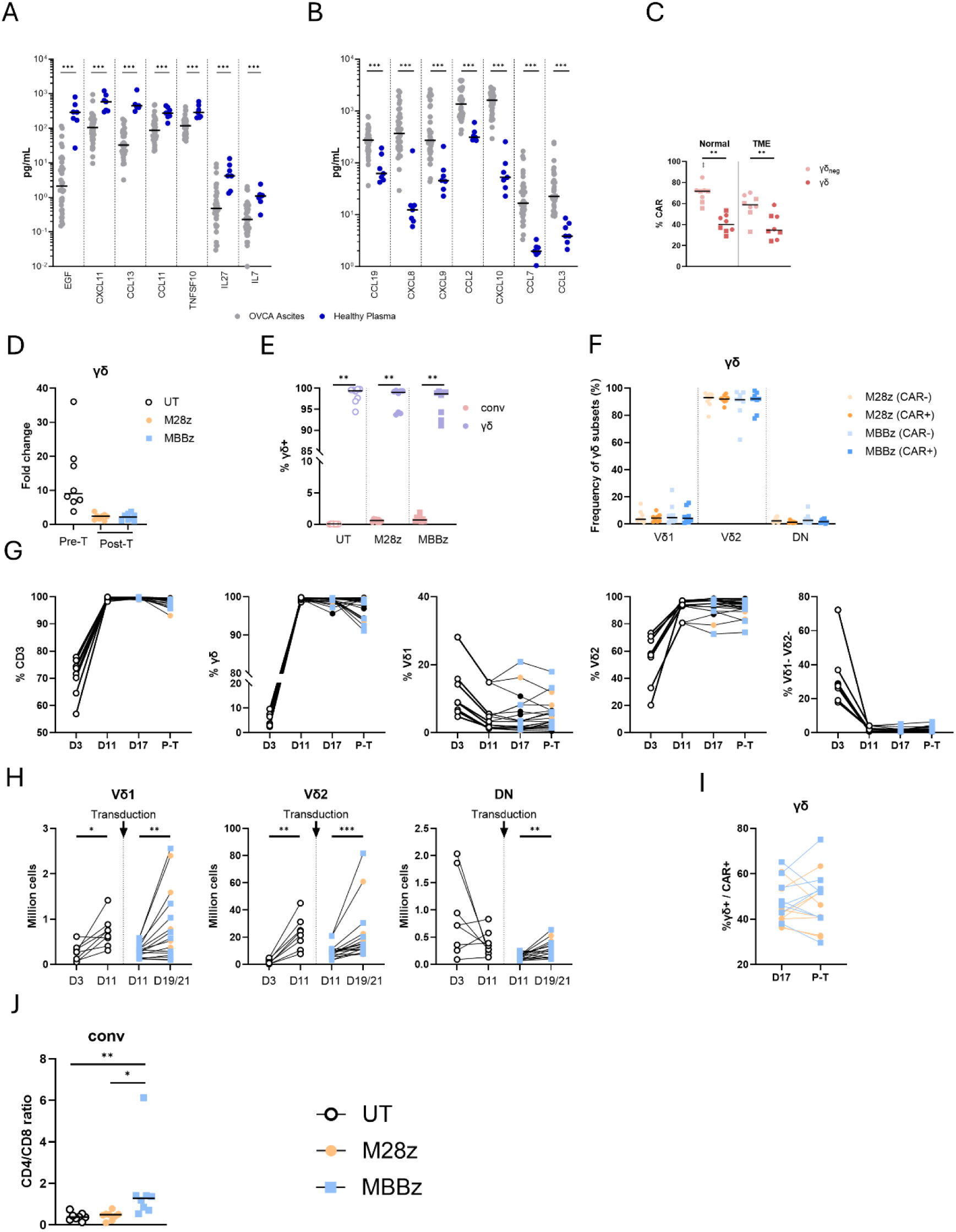
**A-B.** Comparison of cytokine concentration between healthy plasma and OVCA ascites. **C.** CAR (EGFRt+) frequency in γδ- and γδ+ convCAR T-cells, after 6 days exposure to normal or TME-like conditions. Circles represent M28z- and squares represent MBBz-CAR T-cells. **D.** Fold change of γδCAR T-cells expansion before (Pre-T) and after (Post-T) transduction with M28z or MBBz CAR constructs **E.** Frequency of CD3+ γδ+ cells within γδCAR T-cell and convCAR T-cell products and UT T-cells post freezing-thawing. **F.** Frequency of Vδ1, Vδ2 and double negative (DN, Vδ1-Vδ2-) cells among the CAR+ and CAR_neg_ fraction of γδCAR T-cell products (M28z or MBBz). **G.** Frequencies of CD3+, γδ+ (of CD3+), Vδ1+, Vδ2+, Vδ1-Vδ2-cells in M28z- and MBBz-γδCAR T-cell products throughout production (P-T = post-thawing). **H.** Cell count of Vδ1+, Vδ2+, DN (Vδ1-Vδ2-) cells during the production, i.e. before transduction (D3-D11) and after transduction (D11-D19/21). **I**. CAR frequency in M28z- and MBBz-γδCAR T-cells, pre-(D17) and post-cryopreservation. **J**. CD4/CD8 ratio (among CD3+ γδ_neg_ T-cells) in convCAR T-cell products (M28z and MBBz) and UT T-cells. (**A, B, E, H, I)** Wilcoxon test. **(F, J)** Friedman test with Dunn’s multiple comparisons.

**Figure S2.**
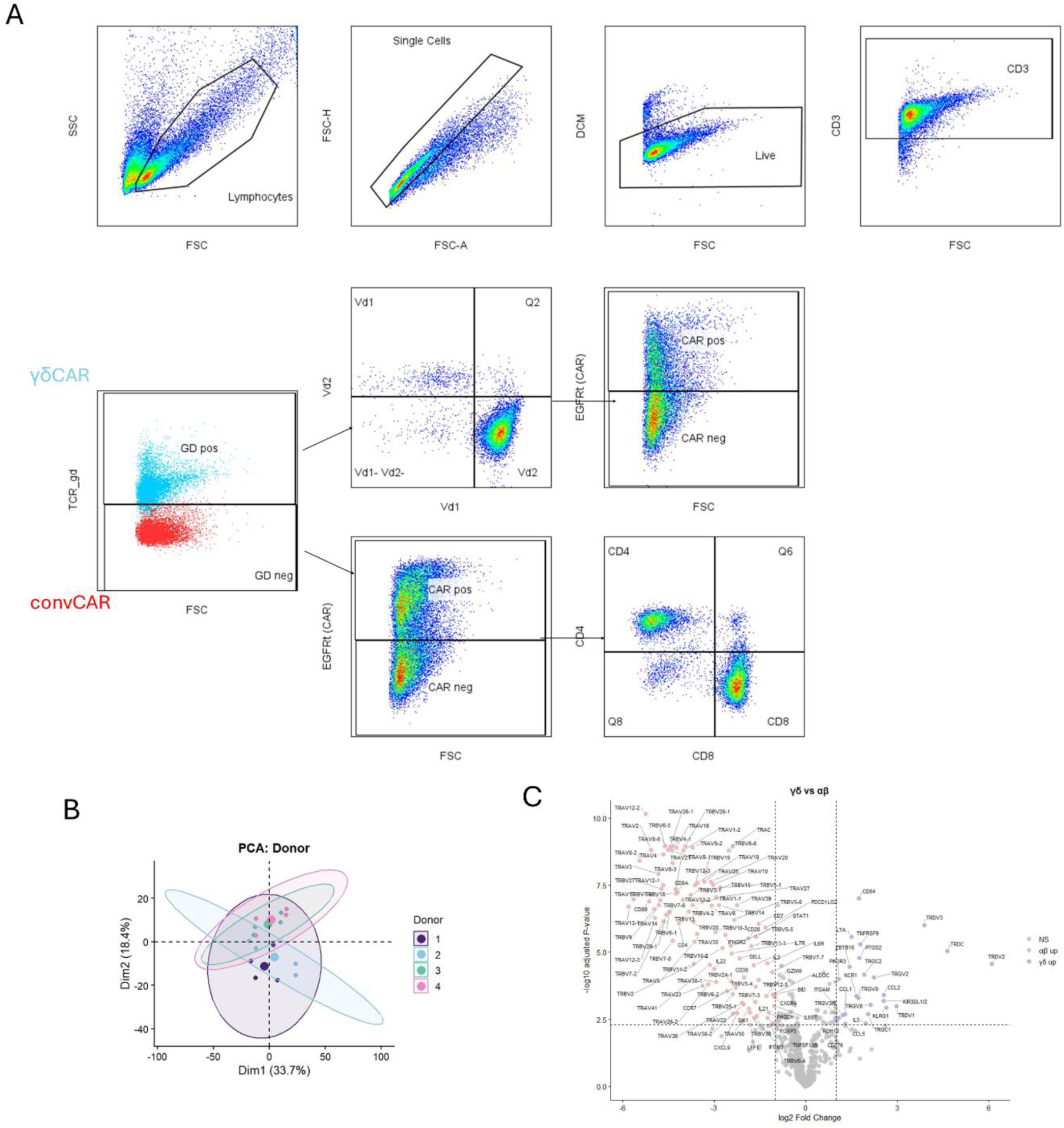
**A.** Flow cytometry gating strategy for CAR T-cells and subsets. **B.** PCA analysis of gene expression profiles comparing 4 donors used for CAR T-cell production. Ellipses represent the 95% confidence interval for each group. **C.** Volcano plot showing differential gene expression between αβCAR and γδCAR T-cells estimated using limma linear modeling with donor and CAR construct correction, with labels for all genes. Adjusted P-value<0.005, |log2FC|>1.

**Figure S3.**
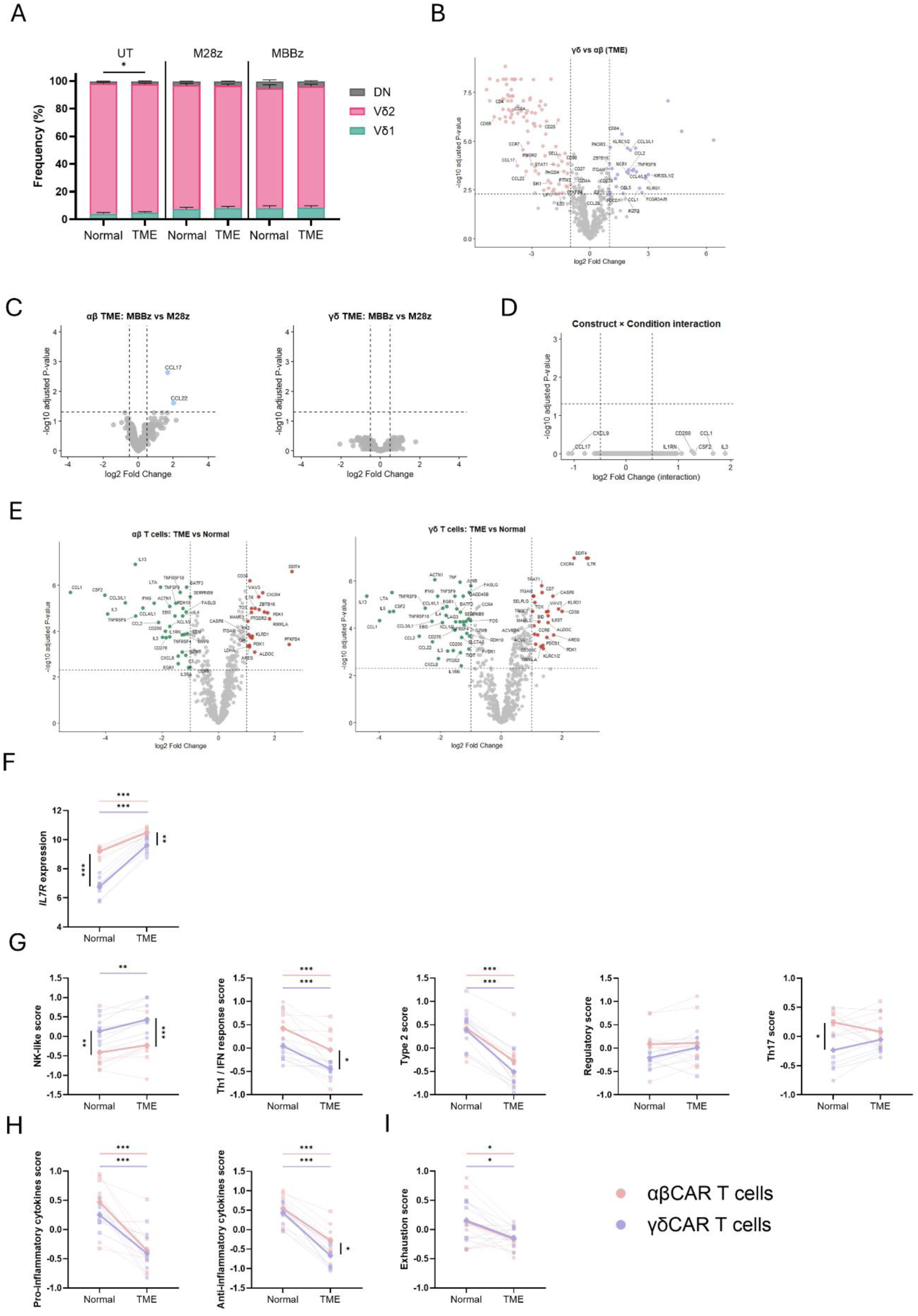
**A.** Frequency of Vδ1, Vδ2 and DN γδ T-cell subsets among UT and M28z- and MBBz-γδCAR T-cells in normal and TME-like conditions. **B.** Volcano plot showing differential gene expression between αβCAR and γδCAR T-cells in TME-like conditions estimated using limma linear modeling with donor and CAR construct correction, with labels for non-TCR genes. Adjusted P-value<0.005, |log2FC|>1. **C.** Volcano plots showing differential gene expression analysis between M28z- and MBBz-CAR T-cell constructs within αβ and γδ CAR T-cells in TME-like conditions estimated using limma linear modeling with donor correction. Adjusted P-value<0.05, |log2FC|>0.5. **D.** Construct × condition interaction. Limma linear modeling with adjustment for cell type and donor was used to identify genes with differential TME responses between M28z- and MBBz-CAR T-cells (adjusted P-value<0.05, |log2 FC|>1). **E.** Volcano plots showing differential gene expression analysis between normal and TME-like conditions in αβ and γδ CAR T-cells estimated using limma linear modeling with donor and CAR construct correction. Adjusted P-value cutoff 0.005, |log2FC| > 1. **F.** Most differentially expressed gene identified by cell type × condition interaction analysis (*IL7R*). Statistical comparisons were performed using donor-paired limma linear models adjusted for CAR construct (n=8 paired observations per comparison). P-values were adjusted across the 664 tested genes using the BH FDR method. **G-I**. Gene signature scores across normal and TME-like conditions. Signature scores were calculated as the average Z-normalized expression of the constituent genes. Statistical comparisons were performed using donor-paired limma linear models adjusted for CAR construct (n=8 paired observations per comparison). P-values were adjusted across the 14 tested modules using the BH FDR method. **(F-I)** Thin lines connect donor-matched samples between normal and TME-like conditions with αβ and γδ T-cells analyzed separately, thick lines indicate mean expression values, and symbols denote construct identity (circles: M28z; squares: MBBz, diamonds: group means). Significance is shown as adjusted P-values (q-values): *q < 0.05, **q < 0.01, ***q < 0.001.

**Figure S4.**
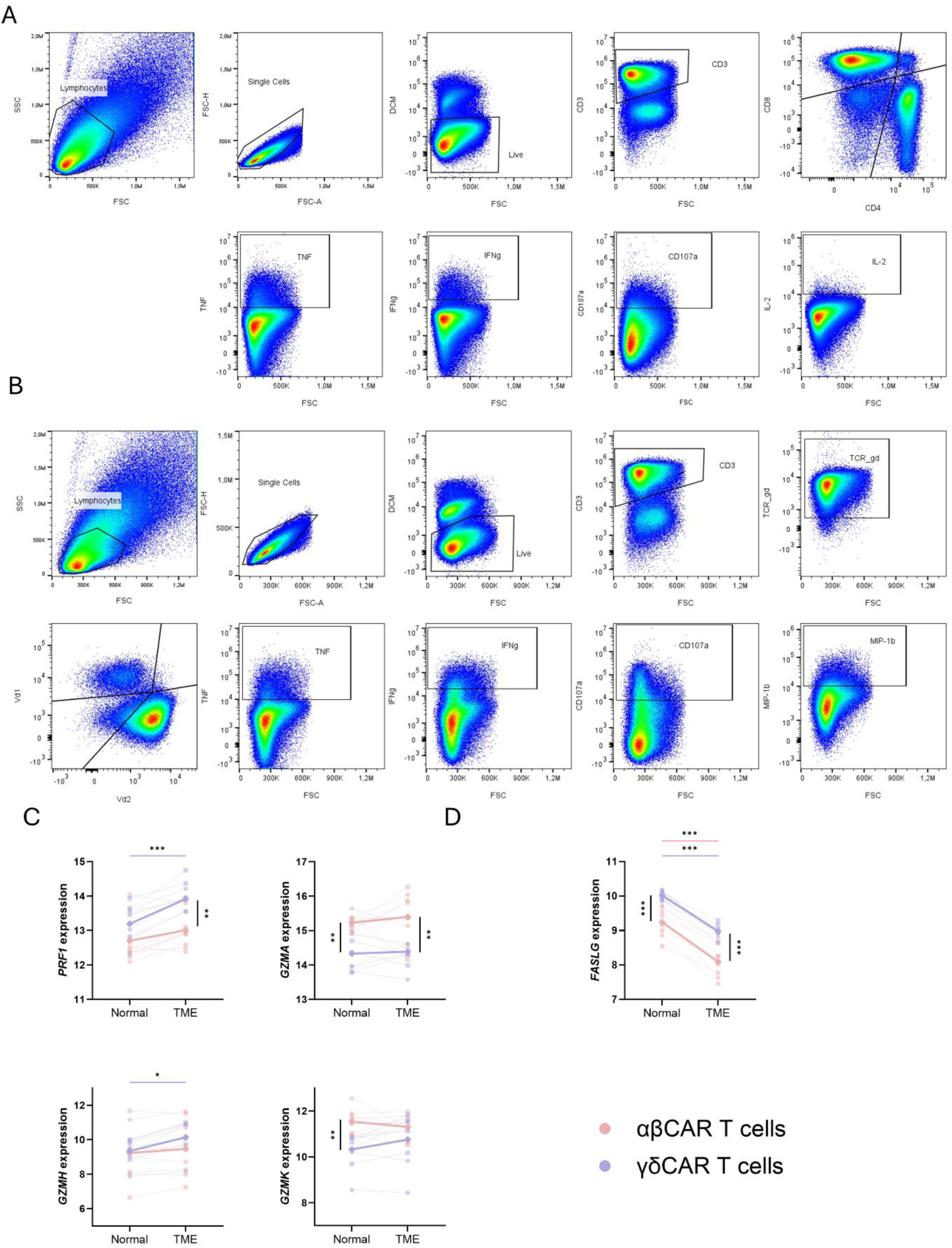
**A.** Gating strategy of functional assay in convCAR and UT T-cells. **B.** Gating strategy of functional assay with intracellular cytokine staining in γδCAR and UT T-cells. **C-D.** Individual gene expression across normal and TME-like conditions in αβCAR and γδCAR T-cells. Thin lines connect donor-matched samples between normal and TME-like conditions with αβ and γδ T-cells analyzed separately. Thick lines indicate mean expression values, and symbols denote construct identity (circles: M28z; squares: MBBz, diamonds: group means). Statistical comparisons were performed using donor-paired limma linear models adjusted for CAR construct (4 donors, 8 donor–construct paired observations per comparison). P-values were adjusted across the 664 tested genes using the BH FDR method. Significance is shown as adjusted P-values (q-values): *q < 0.05, **q < 0.01, ***q < 0.001.

**Figure S5.**
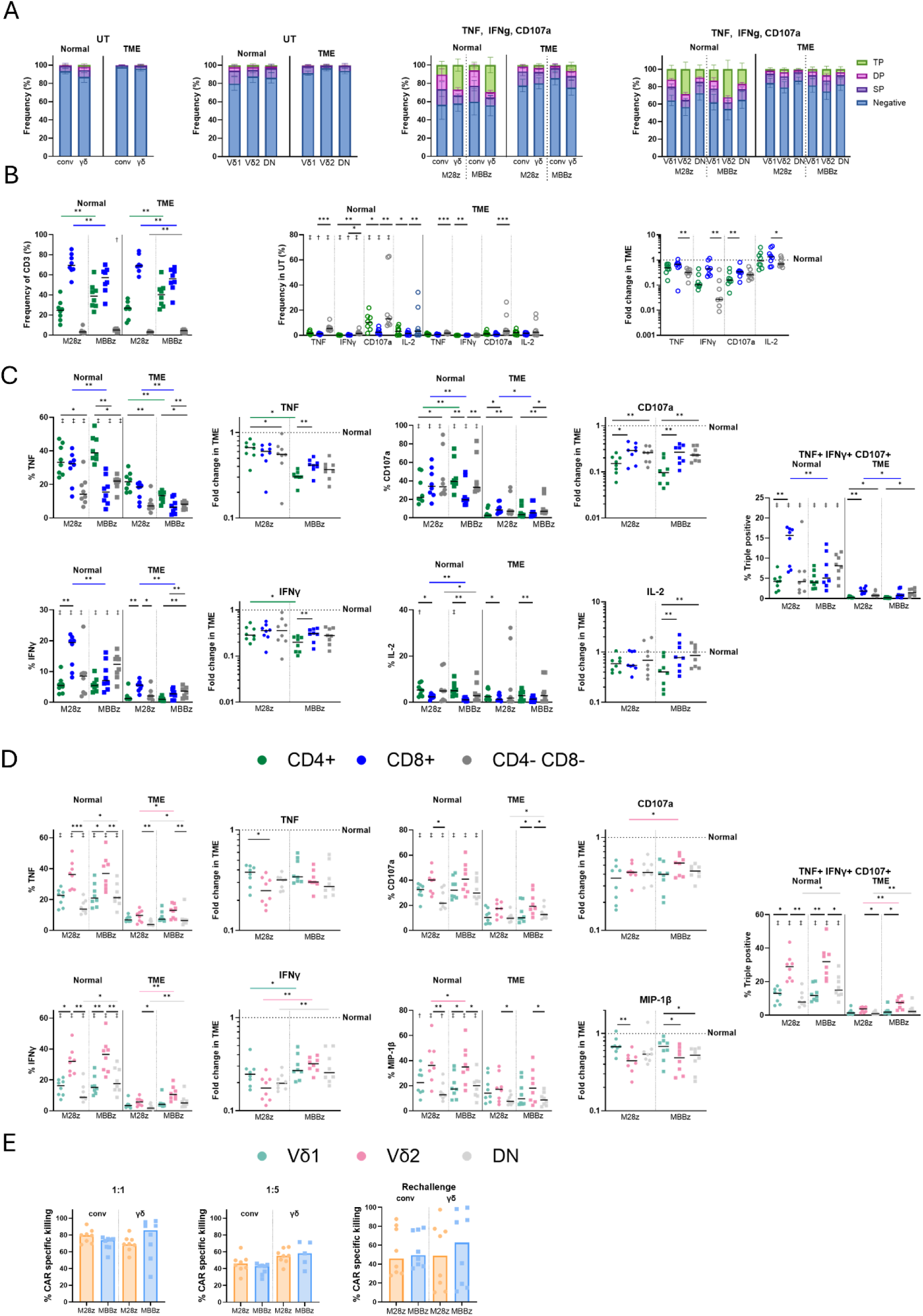
**A.** Frequency of TNF, IFNγ and CD107a triple negative, single positive, double positive, and triple positive cells in UT conv T-cells, γδ T-cells and γδ T-cell subsets, and in the corresponding M28z- and MBBz-convCAR T-cells, γδCAR and and γδCAR T-cell subsets. **B.** Frequency of CD4+, CD8+ and CD4- and CD8-subsets in convCAR T-cells, and frequency of TNF+, IFNγ+, CD107a+ and IL2+ cells in those subsets of UT conv T-cells. **C.** Frequency of TNF+, IFNγ+, CD107a+ and IL2+ cells in CD4+, CD8+ and CD4- and CD8-subsets of M28z- and MBBz-CAR T-cells in normal and in TME-like conditions, accompanied by corresponding fold change of TNF+, IFNγ+ and CD107a+ frequency between TME-like and normal conditions. **D.** Frequency of TNF+, IFNγ+ and CD107a+ cells in Vδ1, Vδ2 and DN subsets of γδCAR T-cells in normal and in TME-like conditions, followed by fold change of TNF+, IFNγ+ and CD107a+ frequency between TME-like and normal conditions and frequency of triple-positive TNF+ IFNγ+ CD107a+ cells in normal and in TME-like conditions. **E.** CAR-specific cytotoxicity of γδCAR T-cells and convCAR T-cells in TME-like conditions after 24h-stimulation with OVCAR3 cells at 1:1 (left) and 1:5 (middle) E:T ratio, and after 3 repeated stimulations with OVCAR3 cells at 1:1 ratio over 6 days (with cytotoxicity measurement 24h after the third stimulation (right). **(C-D)** Wilcoxon test was performed for paired comparisons constructs, cell subsets or cell types, (represented by stars) and conditions (represented by daggers) */† P<0.05, **/‡ P<0.01. TP: triple positive, DP: double positive, SP: single positive.

**Figure S6.**
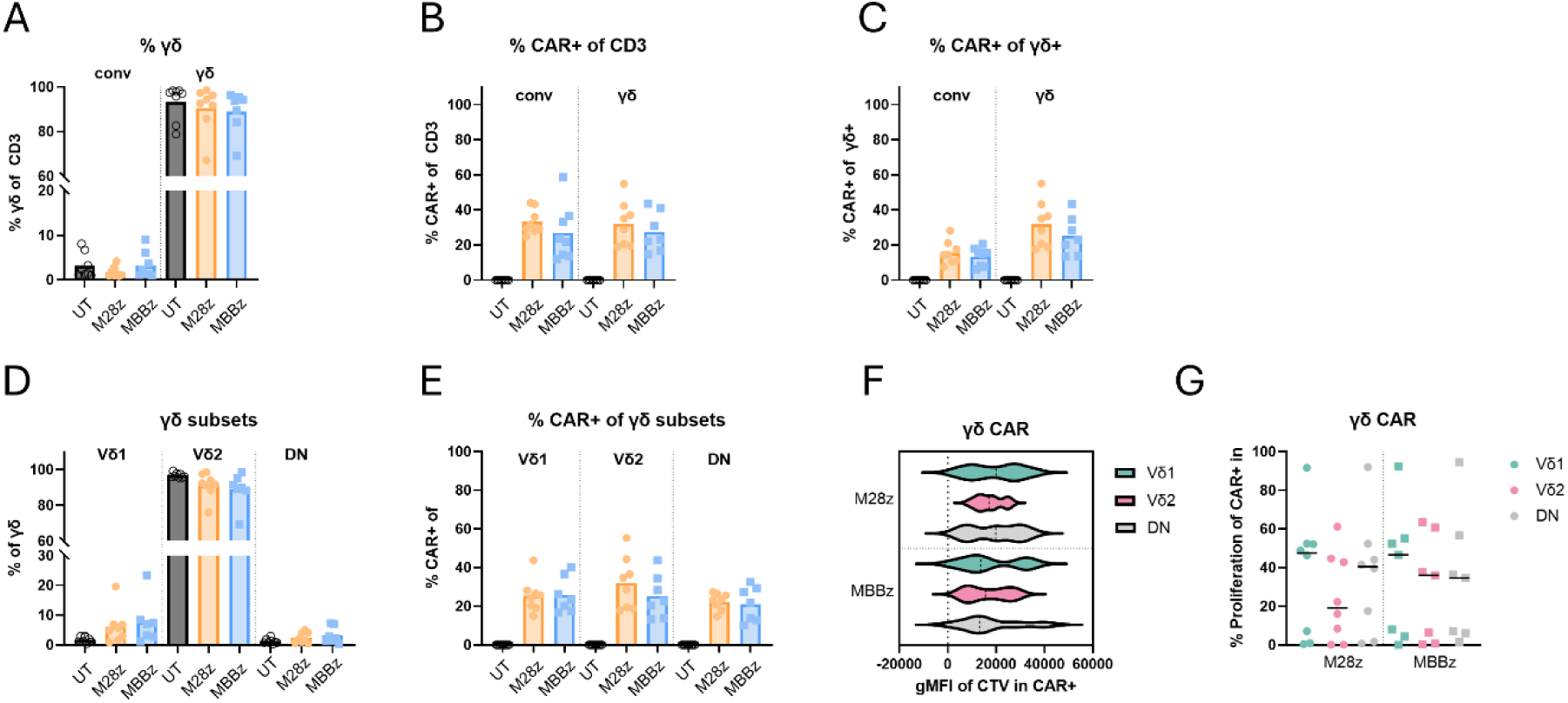
Cell content characterization after the proliferation assay. **A.** Frequency of γδ+ cells in convCART-cells and γδCAR T-cells. **B-C.** Frequency of CAR+ cells among CD3 **(B)** and among γδ **(C)** in convCART-cells and γδCAR T-cells. **D.** Frequency of γδ T-cell subsets (Vδ1, Vδ2 and DN) in γδCAR T-cells and UT γδ T-cells. **E.** Frequency of CAR+ cells in each γδ T-cell subset (Vδ1, Vδ2 and DN) in γδCAR T-cells and UT γδ T-cells. **F-G.** Proliferation of γδCART-cell subsets (Vδ1, Vδ2 and DN) as gMFI of CTV in the CTV-fraction among CAR+ cells (**F**) and frequency of CTV in the CTV-fraction of CAR+ cells **(G). (F,G)** Friedman’s test with Dunn’s multiple comparisons was used for paired comparisons between cell subsets and Wilcoxon test was used for paired comparisons between constructs.

**Figure S7.**
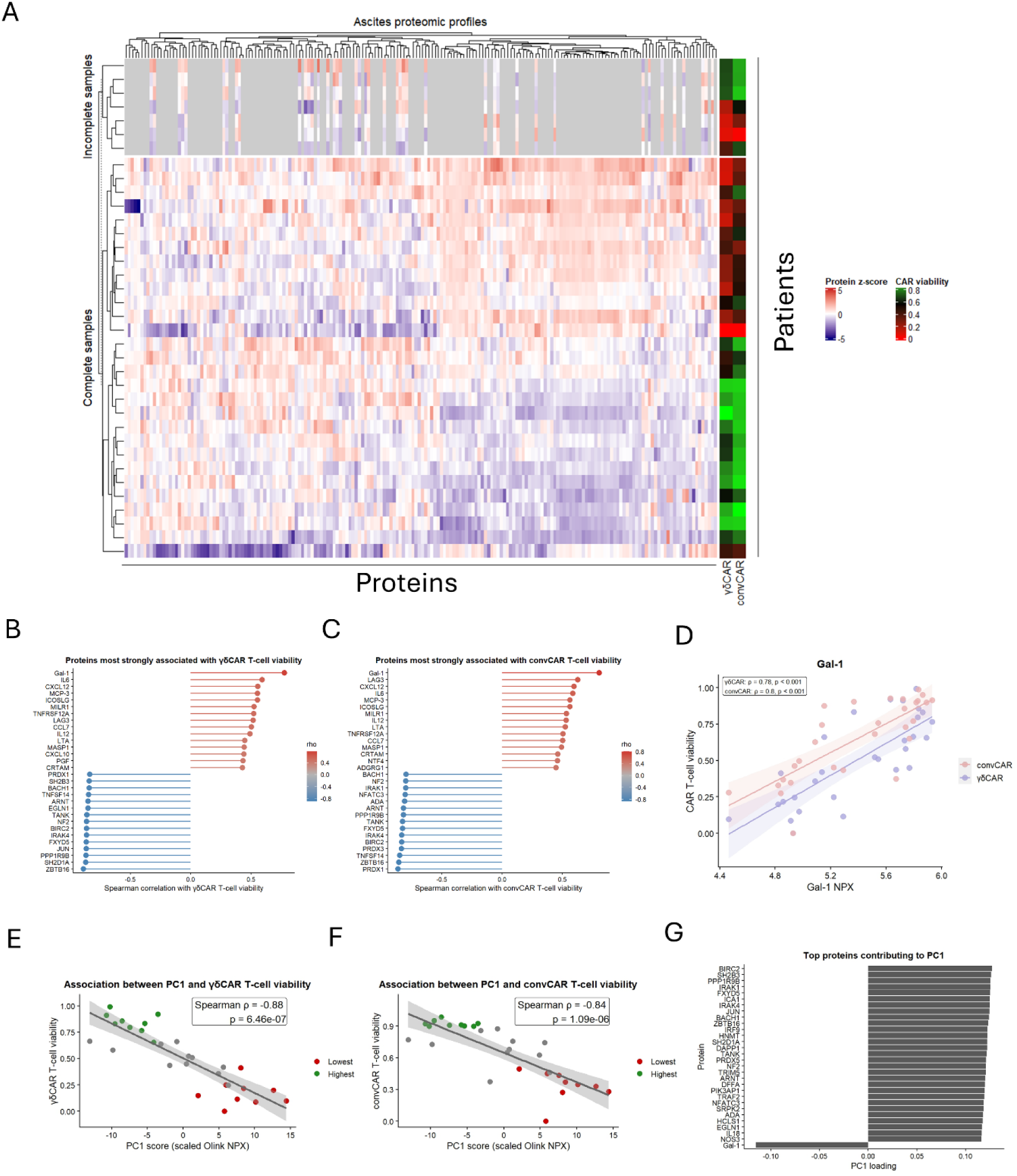
**A.** Heatmap showing protein expression across patient-derived ascites samples. Protein expression values were standardized to protein-wise z-scores across samples, with red indicating above-average and blue indicating below-average protein abundance. Hierarchical clustering of patients and proteins was performed using Euclidean distance and complete linkage, with the clustering structure based on the 29 samples with complete proteomic measurements. Missing protein measurements are shown in grey and respective patients are excluded from protein clustering. γδCAR and conventional CAR T-cell viability are shown as row annotations. **B-C.** Lollipop plots showing proteins with the strongest positive and negative associations with γδCAR T-cell viability **(B)** and convCAR T-cell viability **(C)**. Proteins are ranked according to Spearman correlation coefficients (ρ). Red indicates positive associations and blue indicates negative associations. **D.** Correlation between ascites galectin-1 (Gal-1) abundance and γδCAR and convCART-cell viability. **E-F.** Correlation between the PC1 score from scaled ascites proteomic profiles and γδCAR T-cell viability **(E)** and convCAR T-cell viability **(F)**. **(H)** Top 30 proteins contributing to PC1 variation in ascites proteomic profiles. Proteins are ranked according to the absolute magnitude of their PC1 loading. **(D-F)** Spearman rank correlation was used to assess associations. The lines represent linear regression with 95% confidence interval.

### Data File Legends

Data file S1. Individual-level data

Data file S2. Nanostring data (gene expression) and scores

Data file S3. Olink data (Protein expression)

Data file S4. Gene Ontology analysis

